# Neural categorisation of rhythm in human newborns

**DOI:** 10.64898/2026.10.16.759621

**Authors:** Francesca M. Barbero, Tomas Lenc, Alban Gallard, Nori Jacoby, Rainer Polak, Manuel Varlet, Arthur Foulon, Sahar Moghimi, Sylvie Nozaradan

## Abstract

Music is ubiquitous across cultures, and humans engage in social interactions through music from early in life. A cornerstone of these interactions is rhythm categorisation, allowing the sheer diversity of rhythmic inputs to be mapped onto internal rhythm categories. The pervasiveness of certain rhythm categories across cultures suggests that rhythm categorisation may be partially rooted in neurobiological predispositions constraining the internal representations of rhythm. However, this view appears to conflict with the diversity of music practices across cultures, which points to a role of experience in rhythm categorisation. To assess the relative contributions of neurobiological predispositions and experience, we tested 0- to 3-day-old newborns, capitalising on their minimal postnatal experience with music. Using electroencephalography (EEG), we recorded, in newborns and adults, brain responses to drum sounds whose pattern of relative interval duration ranged from isochronous to long-short rhythms (1:1 – 2:1 interval ratio). Results indicate significant categorisation in newborns’ neural responses, with a representational structure that correlate with those of adults, yet with a greater degree of differentiation across rhythms. These findings reveal neurobiological predispositions enabling rhythm categorisation to be present at birth and further shaped across the lifespan, highlighting the evolutionary roots and plasticity supporting this human cultural practice.

## Introduction

Music is ubiquitous across human cultures (Blacking, 1973; Mehr et al., 2019; Merriam, 1964; Nettl, 1996; Trainor & Hannon, 2013), with engagement in social interactions through music appearing early in life (Aubinet, 2024; Blacking, 1967; Nguyen et al., 2023; Trehub et al., 1993; Young & Ilari, 2019). A cornerstone of these interactions is the capacity to categorise musical rhythms (Trevarthen, 1999). This skill consists in mapping the sheer diversity of possible rhythmic sensory inputs onto a finite set of internal categories of rhythms (Barbero, Lenc, et al. 2025; Clarke, 1987; Desain & Honing, 2003; Jacoby et al., 2024; Jacoby & McDermott, 2017; Polak et al., 2018; Povel, 1981; Ravignani et al., 2016; Repp et al., 2011, 2012; Schulze, 1989; Summers et al., 1986).

Categorisation can be considered a fundamental computational process that our central nervous system needs to implement. Rather than treating each sensory event as singular – a process that would prove highly taxing for the human brain – the brain generalises across sensory inputs from the same category, while differentiating inputs from distinct categories (Fabre-Thorpe, 2011; Goldstone, 1998; Grill-Spector & Weiner, 2014; Holt & Lotto, 2010; Ley et al., 2014). By extension, categorisation makes perception more reliably shareable across individuals and thus opens it to the social basis of perceptual learning in human communication systems, specifically in language acquisition and music practice (Kuhl, 2004, 2011; Pierrehumbert, 2003; Polak et al., 2026; Zaslavsky et al., 2018). In the context of musical rhythm, the ability to map rhythmic sensory inputs onto a set of shared internal rhythm categories is essential for successful social interaction through music, by enabling the species-typical capacity for temporal coordination across individuals, which arguably represents a core function of human rhythm processing (Clayton, 2019; Clayton et al., 2020; Cummins, 2009; Jacoby et al., 2021; Phillips-Silver et al., 2010).

What are the brain processes underlying rhythm categorisation? One view posits that rhythm categories might be at least partially rooted in general properties of neural assemblies (not necessarily unique to humans; De Gregorio et al., 2021, 2024; Roeske et al., 2020). This view has been corroborated by recent observation that certain rhythm categories are pervasive across human cultures (Jacoby et al., 2024). These rhythm categories are characterised by relative interval durations corresponding to ratios of small integers such as equal time intervals (1:1 ratio) and the grouping of equal intervals in twos (2:1 ratio; Fig. 1A).

**Figure 1.**
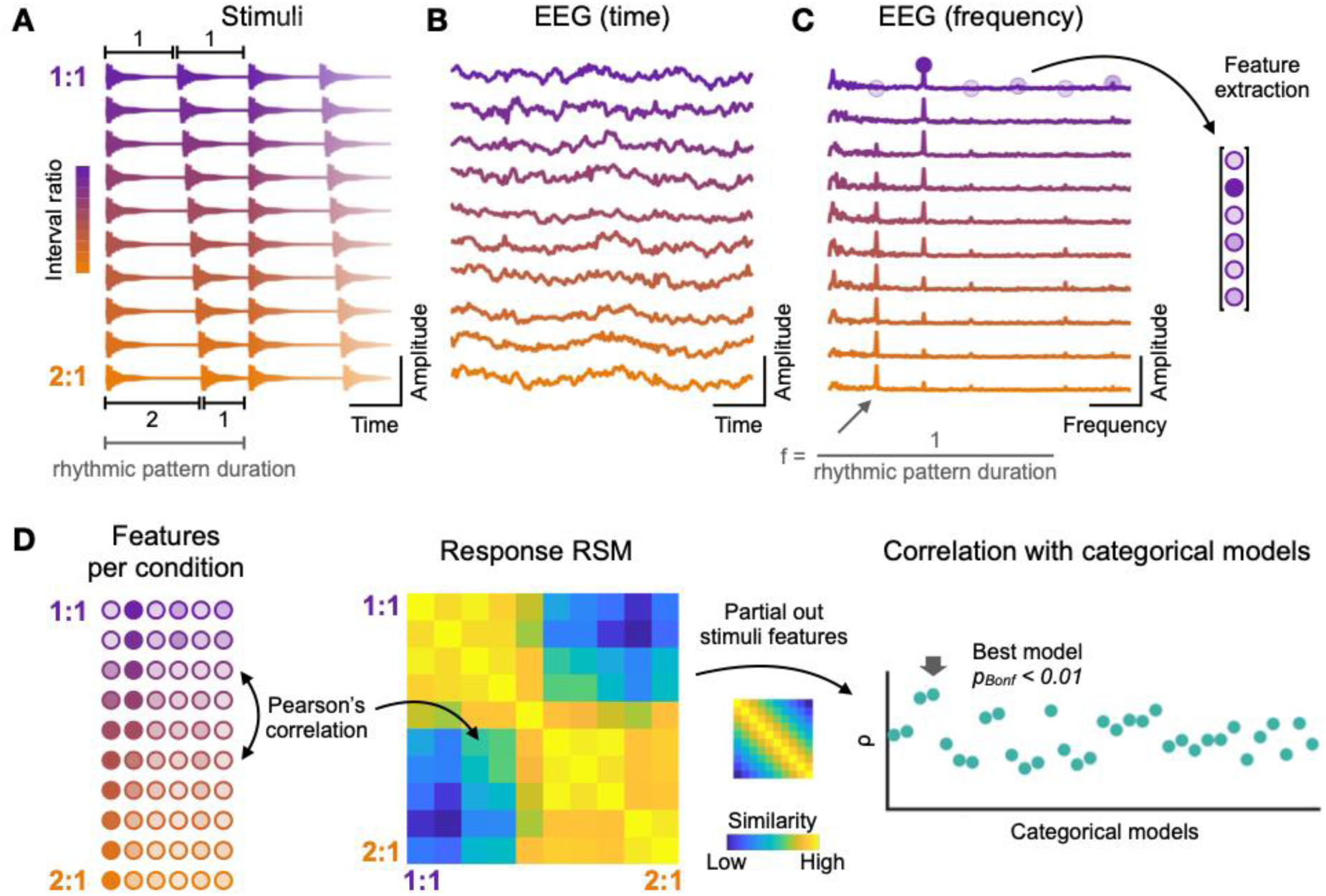
Experimental design and fRSA approach. (A) Stimuli consisted of two-interval rhythmic patterns which were looped over time to obtain 30-s long rhythmic sequences (1.5 s excerpts in the figure, corresponding to 2 repetitions of each rhythmic pattern). The position of the second note of the rhythmic pattern was gradually shifted in time across conditions, yielding a total of 10 conditions with interval ratios spanning from 1:1 (isochrony, top) to 2:1 (long-short pattern, bottom). (B) Averaged time domain EEG responses to each rhythmic sequence (in adults, visualisation purposes). (C) Averaged magnitude spectra of the EEG responses to each rhythmic sequence (in adults, visualisation purposes). Responses exhibited peaks of activity at frequencies corresponding to the repetition rate of the rhythmic pattern and harmonics. The amplitude of the response at these a priori defined frequencies was used to build a feature vector for each condition. (D) A response representational similarity matrix (RSM, middle) was built by correlating feature vectors (left) across all pairs of conditions. Rhythm categorisation was assessed by computing partial correlations between the response RSM and theoretical categorical models while controlling for the contribution of stimulus features. The best categorical model was selected as the theoretical model with the highest correlation coefficient (ρ) and the significance of its fit was obtained using permutation testing (right; p-value Bonferroni corrected for multiple comparisons).

However, the perceptual bias toward these small-integer rhythm categories could also result from perceptual learning and neural plasticity, in face of the pervasiveness of these rhythms in the music-cultural environment. The key role of learning and plasticity in rhythm categorisation is demonstrated by growing evidence for culture-specific rhythm categories, in addition to the ubiquitous small-integer ratio rhythms, that mirror culture-specific features found in the corresponding music-cultural practices and repertoires (Brailoiu, 1951; Clayton, 2020; Cler, 1994; Goldberg, 2016, 2019; Johansson, 2017; Kvifte, 2007; Polak, 2025). Evidence for the role of learning and plasticity in rhythm processing come from cross-cultural studies (Jacoby et al., 2024; Jacoby & McDermott, 2017; Polak, 2025; Polak et al., 2018, 2026; Snyder et al., 2024; Ullal-Gupta et al., 2014) and developmental research (Einarson & Trainor, 2016; Hannon, 2009; Hannon et al., 2011, 2018; Hannon, Soley, et al., 2012; Hannon, Vanden Bosch der Nederlanden, et al., 2012; Hannon & Trainor, 2007; Hannon & Trehub, 2005; Lynch et al., 1990; Snyder et al., 2024; Soley & Hannon, 2010; Trainor & Hannon, 2013). For instance, while 6- to 7-months old infants can detect structure violations in rhythmic patterns from familiar and unfamiliar musical systems (Hannon & Trehub, 2005a), 12-month-old infants show a bias for culturally familiar rhythms, as adults (Hannon & Trehub, 2005b). Altogether, these studies suggest that rhythm categorisation is a plastic function that is shaped by cultural experience across the lifespan (Polak et al., 2026).

Since decades, teasing apart the respective contributions of neurobiological predispositions and experience, as well as their interactions across development, has remained a major methodological challenge for the social, cognitive, and life sciences (e.g., see Lewontin, 2000; Meaney, 2010; Moore, 2003; Tabery, 2014; Turkheimer, 2000). One approach consists of probing perceptual abilities early in life, capitalising on the minimal experience with cultural environments beyond the one developed in utero (Perani et al., 2010). For example, speech processing abilities have been extensively studied in newborns and during the first months of life (Dehaene-Lambertz & Gliga, 2004; Gervain & Mehler, 2010), showing categorical perception of phonemic contrasts (Eimas et al., 1971) that is progressively attuned to the statistics of the native language across the first year of life (Kuhl et al., 1992; Werker & Tees, 1984). Moreover, mismatch responses compatible with phoneme categorisation have been observed from scalp EEG in newborns (Dehaene-Lambertz & Pena, 2001) and in three-month-old infants (Dehaene-Lambertz & Baillet, 1998). However, in the domain of rhythm perception, evidence for categorisation in young infants has been largely restricted to behavioural paradigms until today – paradigms that are either (i) indirect and not readily comparable to adult measures (e.g., preferential looking), or (ii) contingent on sensory–motor reproduction abilities with protracted developmental trajectory (Nave, 2024), thus not applicable to early childhood and infant cohorts.

Here we advance beyond this gating bottleneck by providing evidence for rhythm categorisation in human newborns, leveraging a methodological approach that we recently developed to capture rhythm categorisation directly from neural activity, without relying on any behavioural measure (Barbero, Lenc, et al. 2025). Neural responses were recorded using high density electroencephalography (EEG) for optimal measurement reliability. In addition, neural and behavioural responses to the same rhythms were recorded separately in adult participants, allowing direct comparison of the categorical structures emerging across different ages and measurements. We found that newborns’ and adults’ neural responses did not merely track the temporal structure of the stimuli, but also exhibited significant rhythm categorisation. Moreover, while the geometry of newborns’ rhythm representation correlated with the ones observed in adults, it also showed a greater degree of differentiation across rhythms as compared to adults.

Our results provide evidence for neurobiological predispositions that may bias the representation of rhythmic inputs at birth and serve as a scaffold for further tuning of rhythm categorisation through experience across the lifespan. Crucially, the present study lays the groundwork for future research aiming at elucidating the early developmental trajectory of rhythm categorisation, how it is shaped by cultural experience, and how this function may depend on a network of brain regions possibly shared in part with nonhuman species.

## Results

We used scalp electroencephalography (EEG) to measure brain responses to auditory rhythmic sequences in a group of healthy, full-term, newborn participants (N = 38, 20 females, age range at testing: 0-3 days). Participants were presented with various rhythmic sequences, each made of a repeated rhythmic pattern. Each pattern consisted of two identical percussive sounds whose onsets delimited two-time intervals. The range of relative durations of these two intervals was defined by two extreme values: a 1:1 ratio (i.e., the two time intervals have the same duration: isochrony) and a 2:1 ratio (i.e., the first interval is twice as long as the second one, creating a long-short pattern; Fig. 1A, Supplementary table 1). These two ratios were then morphed to obtain eight additional time intervals ratios evenly covering the range between the 1:1 and 2:1 ratios, thus yielding a continuum of rhythmic ratios across 10 conditions in total (Fig. 1A). This continuum of rhythmic stimuli was selected based on a number of studies showing that adult Western Europe-enculturated participants perceive rhythms along the 1:1 – 2:1 gradient as two discrete rhythm categories (Barbero et al., 2025; Clarke, 1987; Jacoby & McDermott, 2017; Polak et al., 2018; Povel, 1981; Repp et al., 2012).

Neural responses to two-interval rhythms were analysed using a methodological approach combining frequency-domain (Lenc et al., 2018, 2021; Norcia et al., 2015; Nozaradan et al., 2012, 2018) and representational similarity analyses (RSA; Kriegeskorte, 2008; Kriegeskorte & Kievit, 2013), hereafter referred to as fRSA (Barbero, Lenc, et al. 2025).

### Evidence for neural categorisation of rhythm at birth

Preprocessed neural responses recorded in newborns were analysed in the frequency domain. The advantage of using rhythmic patterns that repeat periodically over the sequences lies in the fact that the elicited brain responses distribute over specific frequency bins determined a priori by the pattern repetition rate (i.e., the repetition frequency and its harmonics) (Norcia et al., 2015; Rossion, 2014). Moreover, how the response is distributed across harmonics reflects the temporal structure of the sensory events composing the pattern (Lenc et al., 2018, 2021, 2024; Nozaradan et al., 2012, 2018).

Following common practices adopted in frequency-tagging analyses (Barbero et al., 2021, 2025; Rekow et al., 2021; Retter, 2016; Volfart et al., 2021), we first determined the range of consecutive harmonics of the pattern repetition frequency with response amplitude significantly higher than noise. To do so, we averaged the time domain data across newborns, conditions, and a set of fronto-central electrodes (Fig. 2A, right panel). The choice of electrodes was based on the observation that neural responses to auditory rhythmic stimuli in infants are best captured by fronto-central electrodes (Cirelli et al., 2016; Edalati et al., 2022, 2023, 2024; Lenc et al., 2023; Saadatmehr et al., 2025). This was confirmed by visual inspection of the topography of the response obtained in the current study (Fig. 2C). The data was then submitted to a Fast Fourier Transform (FFT) to obtain a magnitude spectrum, and the relative prominence of the a priori determined frequencies of interest with respect to local noise baseline was calculated as z-score of their respective magnitude versus their surrounding frequency bins (frequencies of interest: f = 1/rhythmic pattern duration = 1.33 Hz and harmonics; surrounding frequency bins range: ± 0.167 Hz; see Materials and Methods) (Barbero et al., 2021; Retter, 2016; Volfart et al., 2021). Our analysis revealed magnitude significantly above the local noise baseline from the second to the fifth harmonics of the pattern presentation rate (2f = 2*1.33 Hz = 2.66 Hz, and so on; z > 3.09, corresponding to p < 0.001, one-tailed, signal > noise; see also Barbero et al., 2025). However, to account for the fact that the 1:1 rhythm, which consists of events repeating at twice the pattern repetition frequency and thus projects only onto even harmonics of the pattern repetition frequency (at 1/0.375 s = 2.66 Hz and harmonics, i.e., 2f, 4f, and so on; see Materials and Methods), we included harmonics from the 1^st^ to the 6^th^ one as frequencies of interest (i.e., harmonics up to 8 Hz: 1.33 Hz, 2.66 Hz, 4 Hz, 5.33 Hz, 6.67 Hz and 8 Hz, as in Barbero, Lenc, et al., 2025).

**Figure 2.**
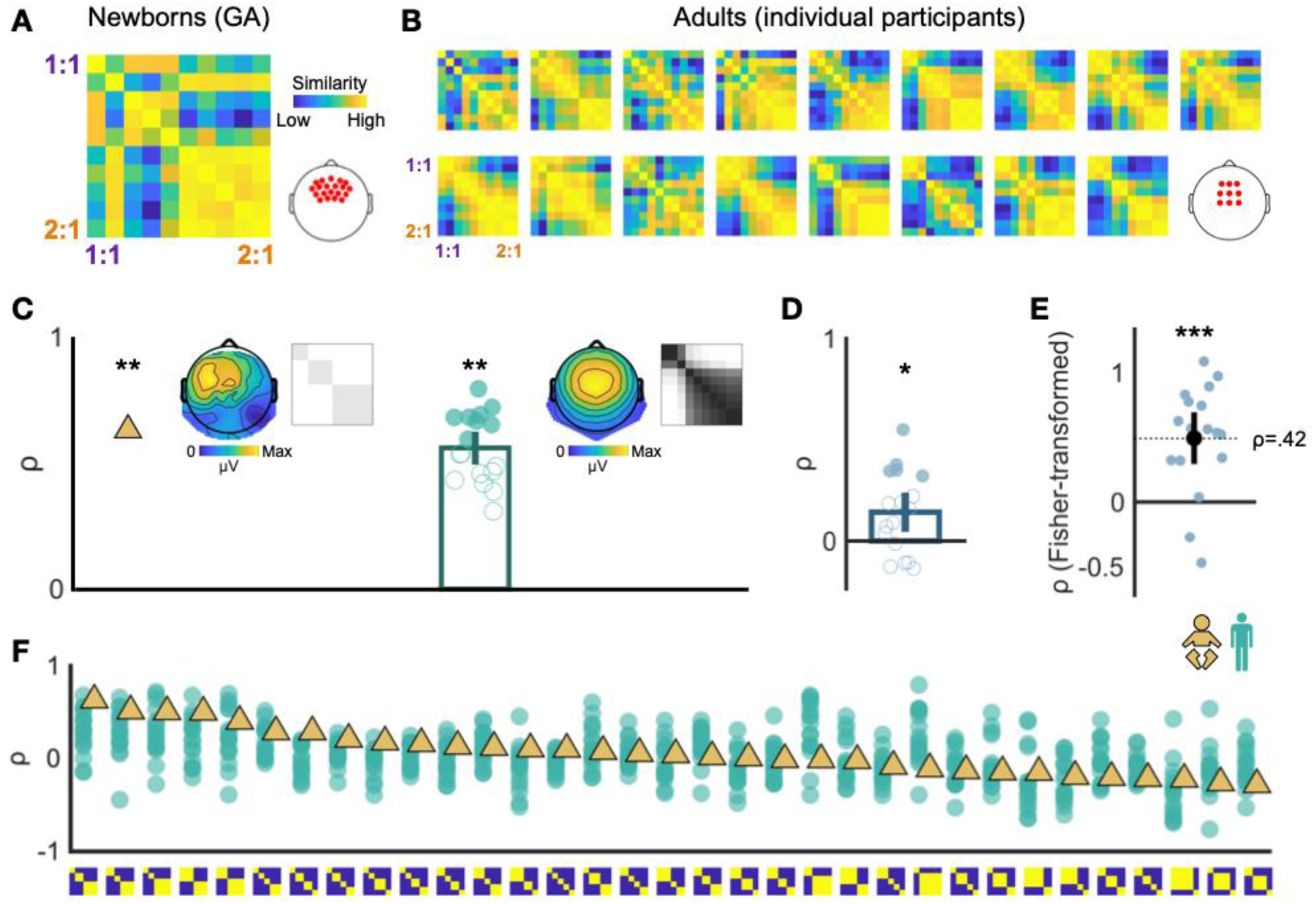
Evidence for neural rhythm categorisation in newborns and in adults. (A) Representational similarity matrix (RSM) obtained from frequency-domain (magnitude) newborn data (group-level average; left). The RSM was obtained from responses at a cluster of fronto-central electrodes (highlighted in red, right). (B) Corresponding RSMs for individual adult participants. (C) Correlation of the neural RSM with the best-fitting theoretical model shown separately for newborns (left) and adults (right). Insets show the best fitting theoretical models (the one identified from the newborns group-level average, and the overlaid ones identified for each adult participant), and the topographical distribution of neural response amplitudes averaged across participants and conditions. (D) Partial correlation of neural RSMs across newborns (grand-average) and adults (individual participants), accounting for the stimulus RSM. (E) Fisher-transformed correlation between the distribution of correlation coefficients across all categorical models between newborns and adults. (F) Distribution of correlation coefficients across all categorical models in newborns (grand-average, gold) and adults (individual participants, green), with categorical models ordered from left to right by decreasing correlation coefficient in newborns. In all bar plots: the error bars represent the 95% CI, asterisks indicate significant results at the group level (* p < 0.05, ** p < 0.01, *** p < 0.001). In C and D, filled circles indicate significant results at the individual level.

Further analyses were performed on newborns showing a response z-score > 1.28 (i.e., corresponding to p < 0.1, one-tailed, signal > noise), yielding a total of 19 newborns showing a reliable response to acoustic sequences over noise baseline (Fig. S1). Crucially, this criterion is orthogonal to the hypothesis tested: as z-scores are calculated on responses averaged across conditions, they are indicative of a neural response to the repeated rhythmic patterns regardless of potential categorisation of rhythm.

Time domain responses of the selected newborn participant were averaged separately by condition and across newborns, which allowed to further reduce the contribution of activity not temporally aligned across newborns within each condition (Strasburger, 1987; see also Supplementary Results corroborating the group-averaged results with group-level statistics on individual neural responses). These responses were then submitted to a Fast Fourier Transform to obtain a magnitude spectrum for each condition. A representational similarity matrix (RSM) was built based on vectors created by concatenating the magnitude of the response at the frequencies of interest for each condition. We then computed pairwise Pearson’s correlation across conditions’ vectors to obtain the neural RSM (Fig. 1, 2A). In order to obtain statistical evidence for rhythm categorisation, we built theoretical models reflecting our operational definition of categorisation: between-categories discrimination (pairwise similarity = 0, minimum 2 categories per model) and within-categories generalisation (pairwise similarity = 1, minimum 2 temporally contiguous conditions per category) (Kriegeskorte, 2008; Rossion et al., 2015). All possible models meeting these criteria for categorisation were included, yielding 33 theoretical categorical models in total, which differed only in the number of categories and in the category boundary locations (i.e., 7 two-categories, 15 three-categories, 10 four-categories and 1 five-categories model; Fig. S2).

We computed partial correlations between the neural RSM and the theoretical models of rhythm categorisation while controlling for the stimulus RSM. The stimulus RSM reflected the temporal structure of the stimuli (i.e., pairwise distance between inter-onset interval ratios; Fig. 1D), thus accounting for any similarity of neural responses across conditions explained by stimulus-driven temporal encoding. The newborns’ neural RSM significantly correlated with a categorical structure (permutation test, ρ = 0.6314, p_Bonf_ = 0.0033), whose shape suggested that rhythmic patterns along the 1:1 – 2:1 temporal continuum are represented as three discrete rhythm categories at birth (Fig. 2C). Moreover, newborns’ neural responses also showed high correlation with a range of models consisting of two or three categories, all characterised by a bigger rhythm category on the side of the 2:1 ratio (i.e., encompassing 5-6 conditions over the 10 conditions, as depicted in Fig. 2F; ρ_model 9_ = 0.6314, ρ_model 13_ = 0.5141, ρ_model 8_ = 0.5055, ρ_model 4_ = 0.5010, ρ_model 3_ = 0.4027 - see also Fig. S2 for larger size visualisation of the theoretical models, Fig. S9, and Supplementary Material for results on different data descriptors).

Lastly, we performed two additional analyses using representational geometries derived from: (i) the acoustic envelope features of the stimuli, and (ii) modelled auditory nerve responses as stimulus RSMs in the partial correlation. Like interval ratios, the acoustic envelope RSM captures the temporal structure of the stimuli, but allows to follow the exact same frequency-specific analysis applied to the neural responses. We extracted the envelope of the acoustic stimuli using Hilbert transform, and submitted the time course to a Fast Fourier Transform. As for the recorded neural responses, the acoustic envelope RSM was built by calculating pairwise Pearson’s correlations between magnitudes at the same frequencies of interest, as performed in the EEG analysis above. The auditory nerve RSM was used to account for potential contributions of early subcortical nonlinearities to rhythm categorisation. To do so, we used a well-established, biologically plausible model which provides simulation of the physiological processes associated with cochlear nonlinearities, inner hair transduction process, the synapse between the hair cell and the auditory nerve, and the associated firing rate adaptation (Bruce et al., 2018). For each rhythmic sequence (i.e., experimental condition), we simulated the time course of the auditory nerve firing rate in response to the stimuli (Fig. S5A). The obtained time courses were submitted to a Fast Fourier Transform, and the auditory nerve RSM was computed by performing pairwise Pearson’s correlations between magnitude responses at the same frequencies of interest (Fig. S5B). The newborns’ neural RSM showed a significant correlation with the same three-category model while partialling out the acoustic envelope RSM (permutation test, ρ = 0.6368, p_Bonf_ = 0.0033) and the auditory nerve RSM (permutation test, ρ = 0.6417, p_Bonf_ = 0.0033), suggesting that rhythm categorisation at birth cannot be merely explained either by the acoustic envelope features of the stimuli or by nonlinear transformations of the stimuli produced at the periphery of the auditory pathway.

### Rhythm representations correlate across newborns and adults

So far, we have shown that rhythmic patterns located along the 1:1 – 2:1 continuum are represented as discrete rhythm categories in the human brain at birth. However, are these categories similar in newborns and adults? To test this, we recorded EEG in a group of healthy adult participants while they listened to the same set of stimuli as did the newborns (N = 19, 13 females, mean age ± SD = 25.57 ± 5.13 years, range: 19.29 – 38.26 years; Western Europe-enculturated sample; 2 adults were excluded from further analyses after preprocessing, see also Materials and Methods). Participants were instructed to listen to the rhythmic sequences without moving, while performing an orthogonal stimulus intensity change detection task to ensure attention was maintained. In separate blocks, participants were asked to tap their finger in synchrony with the rhythmic sequences, as a behavioural measure of rhythm categorisation (see Supplementary Material).

As for the data collected in newborns, pre-processed neural responses from adults were analysed using the fRSA approach (Barbero, Lenc, et al. 2025). These analyses were based on the first 6 harmonics (i.e., up to 8 Hz), after confirming that the magnitude of the responses at these harmonics was significantly above local noise baseline (z > 3.09, p < 0.001, one-tailed signal > noise), thus consistent with the analytical pipeline implemented in newborns and in a previous study probing categorisation of similar rhythmic sequences in adults (Barbero, Lenc, et al. 2025) (see also Supplementary Results for an additional analysis that corroborated the findings with a higher number of harmonics).

We computed the adult neural RSMs by considering magnitudes of responses at the frequencies of interest over a pool of fronto-central electrodes for each participant and condition (Barbero, Lenc, et al. 2025; Lenc et al., 2018, 2020b; Nozaradan et al., 2012; Tierney & Kraus, 2015) (Fig. 2B). We then performed partial correlations between the neural RSMs and the same theoretical models of rhythm categorisation as used for the newborns’ data analysis, while also accounting for the stimulus RSM. Neural responses significantly correlated with theoretical models of rhythm categorisation at both the group and individual level (group-level: permutation test, ρ = 0.5601, p_Bonf_ = 0.0033; individual-level: significant correlation with a theoretical model in 8 out of 17 participants, permutation test, ps_Bonf_ < 0.05; Fig. 2C).

Rhythm categorisation was also observed from behavioural responses recorded during sensorimotor reproduction of the rhythmic stimuli by the same participants in a different session, showing a convergence across behavioural and neural categorisation of rhythm in adults, in line with previous results using fRSA (see Supplementary Material) (Barbero, Lenc, et al. 2025). We also obtained similar results when partialling out the acoustic envelope RSM and the auditory nerve RSM, suggesting that acoustic envelope features and nonlinearities at the auditory periphery cannot explain rhythm categorisation measured with scalp EEG in adults (acoustic envelope: permutation test, ρ = 0.6034, p_Bonf_ = 0.0033, significant results at the individual level in 10 out of 17 participants, ps_Bonf_ < 0.05; auditory nerve responses: permutation test, ρ = 0.5995, p_Bonf_ = 0.0033; significant results at the individual level in 10 out of 17 participants, ps_Bonf_ < 0.05).

To investigate whether we could observe a shared representation of internal rhythm categories between newborns and adults, we compared the fRSA outputs derived from the magnitude spectra of grand-averaged newborn neural data and individual adult neural data (Xie et al., 2022). To this end, we correlated the newborn RSM with each individual adult neural RSM while partialling out the stimulus RSM. Neural RSMs showed a weak but significant correlation across newborns and adults (permutation test, ρ = 0.1423, p = 0.0330, results significant at the individual level in 5 out of the 17 adult participants, ps < 0.05; Fig. 2D). We also correlated the distributions of correlation coefficients across categorical models between the two groups. The distributions significantly correlated across newborns and adults (mean ρ across participants = 0.4156, one-sample t-test against zero after Fisher transformation of the correlation coefficients, p = 0.0002; Fig. 2E, F).

### Different categorical structure of rhythm between newborns and adults

Thus far, our results have revealed (i) significant neural categorisation of rhythm at birth and

(ii) a weak but significant convergence between the neural representations of rhythm recorded with EEG in newborns and adults. As a next step, we investigated potential differences in the format of rhythm categorical structure across the two groups.

First, we tested whether the categorical model that best explained adult neural responses to rhythm could efficiently describe rhythm categorisation also in newborns. The categorical model that best explained adult neural responses (i.e., the model with highest correlation value over the majority of adult participants) was the two-categories model with categorical boundary after condition 3 (model 2, winning model in 6 out of 17 participants). When we performed partial correlation between the newborns’ neural RSM and this categorical model while accounting for the temporal information in the stimuli, results were not significant (permutation test, ρ = −0.0123, p = 0.5319). Inversely, when we performed partial correlations between neural adult RSMs and the categorical model that best explained newborns neural responses (i.e., model 9, three categories, showing the highest correlation value) while partialling out the stimulus RSM, we observed a significant correlation at the group level (permutation test, ρ = 0.2895, p = 0.0053; with results significant at the individual level in 10 out of 17 participants, ps < 0.05). In other words, while the three-category structure that best explained newborns’ responses also accounted for adults’ responses, the opposite was not true: the two-category structure that best explained the adults’ neural responses did not account for the categorical structure of newborns responses.

Second, we implemented a qualitative analysis combining multidimensional scaling and hierarchical clustering to explore the representational structure of rhythm categories in adults and newborns (Mattioni et al., 2020). Specifically, with hierarchical clustering we visualised which rhythm conditions clustered together in brain representations according to a predefined number of clusters. To parallel the number of categories included in the theoretical models of the fRSA analyses, we progressively considered either 2, 3, 4, or 5 clusters (i.e., rhythm categories).

In adults, conditions visually clustered into two categories of rhythms, in line with the results of the fRSA analysis (Fig. 3). This categorical distinction was evident when hierarchical clustering was stopped at 2 clusters, where conditions belonging to the same clusters matched the conditions of the categorical model best explaining adult neural responses (i.e., model 2, clustering together conditions from 1 to 3 versus conditions from 4 to 10). As the number of clusters progressively increased, a third cluster including intermediate conditions appeared (conditions 4, 5 and 6). Lastly, implementing hierarchical clustering with 4 and 5 clusters yield to the formation of clusters including single rhythmic conditions. While theoretical models including rhythm categories made of only one rhythm condition were not implemented in fRSA as they cannot test for within-category generalisation, and as the rhythm conditions forming isolated clusters were not visually segregated from neighbouring conditions, we interpret these results including a high number of clusters as over-partitioning. In other words, these results suggest that clustering the data in 2 or 3 clusters (i.e., rhythm categories) might be sufficient to explain the representational structure of the neural responses in adults, and that including a higher number of clusters might result in overfitting of the neural responses.

**Figure 3.**
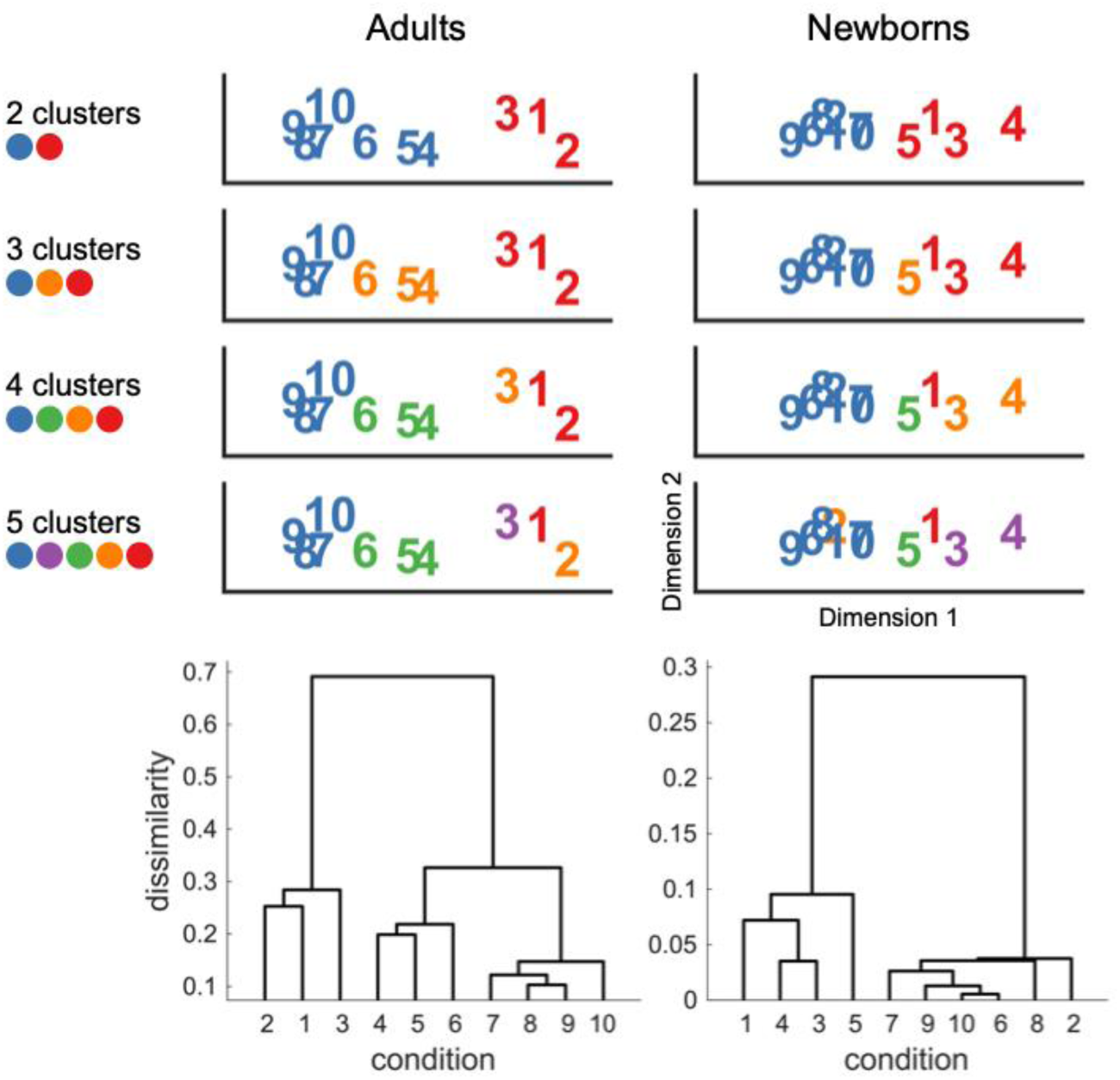
Multidimensional scaling and hierarchical clustering of adults’ and newborns’ neural responses reveal the similarities and differences in the format of rhythm categories across groups. Top: hierarchical clustering was implemented on adults’ (left) and newborns’ (right) RSM considering either 2, 3, 4, or 5 clusters (separate rows). In each panel, conditions belonging to the same cluster are visualised with the same colour. Bottom: dendrograms of adults’ (left) and newborns’ (right) neural responses; the vertical axes represent the dissimilarity distances across conditions. Hierarchical clustering was performed on dissimilarity matrices (1-RSM). In all plots, condition numbers from 1 to 10 corresponds to interval ratios from the morphed continuum between 1:1 (condition 1) and 2:1 (condition 10).

In newborns, conditions closer to the 2:1 ratio clustered together when considering 2, 3, 4, or 5 clusters. On the other hand, conditions closer to the 1:1 ratio were less similarly represented, as shown by their distance and by the progressive split in separate clusters as the number of predefined clusters increased (Fig. 3). These results are in line with the fRSA results, where categorical models showing the highest correlation coefficients consistently show a rhythm category encompassing 5-6 conditions around 2:1.

Altogether, these results suggest that beyond the observed convergences across neural responses to rhythm recorded in newborn and adults, there are also differences in the format of the underlying rhythm categories across groups, in particular with higher degree of differentiation across conditions within the categories closer to 1:1.

## Discussion

By recording high-density scalp electroencephalography (EEG) in healthy full-term human infants at birth, we show that newborns’ brain activity exhibits a representational geometry that is compatible with the presence of underlying rhythm categories. This representational structure is similar to those of adults’ neural responses to the same rhythms, but also shows significant differences in categorical structure. Together, these results reveal the presence of neurobiological predispositions for rhythm categorisation at birth with further shaping of these rhythm categories across the lifespan.

### Neurobiological predispositions to categorise rhythms at birth

Our results reveal that, already at birth, the human brain represents a continuum of stimuli spanning isochronous to long-short rhythms in terms of discrete categories. These findings indicate that ex-utero experience is not necessary to the emergence of rhythm categorisation, which instead may be rooted in predetermined neurobiological mechanisms.

What is the nature of these mechanisms? One potential candidate could be the detection of dissimilarity between successive temporal intervals. Also referred to as anisochrony detection, this process consists in detecting local deviance, i.e., deviance from the directly preceding onset or temporal interval, which in the current study would lead to teasing apart in distinct categories isochronous versus non-isochronous rhythms (Ehrlé & Samson, 2005). While to our knowledge direct evidence for anisochrony detection at birth is lacking, newborns show sensitivity to rhythmic auditory statistical regularities and to their violations (Bianco et al., 2026; Edalati et al., 2022; Háden et al., 2024; Kushnerenko et al., 2013; Moser et al., 2020; Panzani et al., 2023; Winkler et al., 2009). Our findings could therefore reflect an early predisposition for detecting deviations from isochrony (although not necessarily in an adult-like form), supporting categorisation of isochronous versus non-isochronous rhythms at birth.

Beyond the specific case of isochrony (and sensitivity to local deviance from it), another feature of the stimuli that may have potentially driven categorisation in the present study could be the differential degree of temporal unevenness across rhythmic stimuli (i.e., different variance of the temporal intervals composing the rhythms across conditions), yielding differential degree of neural adaptation. In the auditory system, adaptation is usually referred to as suppression of responses to acoustic onsets occurring in rapid succession, that is, separated from previous onsets by short temporal intervals. (Nelken, 2014; Pérez-González & Malmierca, 2014; Willmore & King, 2023). While adaptation is known to occur as early as in the auditory nerve fibres (Pérez-González & Malmierca, 2014), our results showed significant categorical structure even after partialling out the simulated auditory nerve responses to the rhythms in both the newborns’ and adults’ neural responses. This suggests that fast adaptation processes occurring at the auditory periphery cannot explain, alone, rhythm categorisation in the current study.

However, adaptation processes underlying rhythm categorisation in humans could also stem from subcortical auditory neurons of the ascending auditory pathway. Adaptation at the level of these evolutionarily old brain structures has been proposed to contribute to rhythm processing by fostering grouping effects in non-human animals such as rodents (Rajendran et al., 2017). In this context, future research is needed to provide more direct insight into the role of subcortical auditory neurons in rhythm categorisation, for example using a functional localiser to isolate scalp EEG activity originating mainly from cortical versus subcortical auditory sources in response to rhythmic patterns in humans (Lenc et al., 2020a; Nozaradan et al., 2016, 2018).

Slower adaptation produced at later, cortical stages of the auditory processing could also contribute to the neural categorisation of rhythm observed here (Nelken, 2014; Pérez-González & Malmierca, 2014; Rajendran et al., 2017; Ulanovsky et al., 2004). Such non-linear adaptation processes suppressing the response to acoustic onsets when the inter-onset intervals are shorter than specific temporal intervals might, in principle, support categorisation of rhythmic patterns differing in the variance (unevenness) of their constituting temporal intervals. Specifically, this process could have contributed to teasing apart even rhythms from uneven rhythms, as used in the present study. Notably, the most highly ranked theoretical models explaining both adults’ and newborns’ data in the present study consistently showed a category spanning the more uneven rhythms of the tested continuum (i.e., conditions toward the 2:1 ratio), thus in line with this cortical adaptation hypothesis.

Critically, such a cortical adaptation process would be expected to show distinct temporal dynamics in newborns and adults. This could be due, for example, to changes in the oscillatory regimes of neural activity, along with the development of short and long range connections, or to the relatively immature myelination of the brain at birth (Kostović et al., 2021), yielding overall larger conduction times (Daneshvarfard et al., 2020). Indeed, the maturation of the human auditory cortex is protracted over the course of several years (Chen et al., 2023; Moore & Linthicum, 2007; Schnupp et al., 2012), as also reflected in the developmental changes of the electrophysiological auditory responses (Kushnerenko et al., 2007, 2013). While the relationship between conduction time and adaptation time constraints over development is unlikely to be linear, the evidence for an uneven rhythm category that is smaller in newborns than in adults as found here is compatible with cortical adaptation contributing to rhythm categorisation from birth yet with different time constraints in this age group. The exact non-linear function underlying this process remains to be characterised in future research, for example by comparing the obtained categorical structure across different absolute duration of the rhythmic patterns (i.e., different tempos) and different age groups.

More broadly, it remains to be tested whether newborn neural rhythm categorisation generalises beyond deviance-from-regularity detection and adaptation. One way to address this is to measure neural responses to a continuum of non-isochronous rhythms with equalised variance of the component intervals across conditions, thereby controlling for contributions from both anisochrony detection and adaptation (Mares et al., 2025). If categorisation persists under these constraints, it may depend on processes that emerge later in development, potentially through maturation of functional connections between auditory and higher-level associative and motor cortices. This perspective emphasises that multiple mechanisms may support rhythm categorisation in humans, which could ultimately help explain its plasticity.

Finally, our study provides a framework for future research aimed at investigating the extent to which the neural properties underlying rhythm categorisation processes are shared within the human species but also beyond. Animals such as songbirds and non-human primates can exhibit communicative vocalisations characterised by inter-onset intervals biased towards small-integer ratios (De Gregorio et al., 2021; Roeske et al., 2020). Moreover, electrophysiological studies on non-human animal models have reported the existence of temporally selective neurons (Brosch, 2000; Carlson, 2009; Duysens et al., 1996; Elliott et al., 2011; He et al., 1997; Lewicki & Konishi, 1995; Margoliash, 1983; Zhou, 2010) whose function could contribute to the recognition of temporal patterns spanning from tens to hundreds of milliseconds, as required in many forms of animal - including human - communication (Paton & Buonomano, 2018). In this context, the presence of evolutionarily old neurobiological predispositions for the categorisation of time interval might contribute to shaping the development of music and language abilities in humans from the first days of life.

### Rhythm categories are fine-tuned through experience

Our results demonstrate substantial similarities between newborns and adults’ neural representations. Specifically, neural data of both groups showed evidence for significant categorical structure, beyond faithful encoding of the temporal features of the stimuli. Moreover, the geometry of similarities and dissimilarities of the neural responses across conditions was found to correlate significantly across these groups, in particular with a consistent category spanning the more uneven rhythmic stimuli.

However, our results also revealed significant differences across the two groups. Crucially, while a three-category structure best explained newborns’ neural data, three-categorical models were not predominant in adults’ neural or behavioural data. Instead, both neural and behavioural responses were more often best explained by two-category models in adults (with bigger category spanning the uneven rhythms, in line with previous findings; Barbero, Lenc, et al., 2025; Clarke, 1987; Desain & Honing, 2003; Jacoby et al., 2024; Jacoby & McDermott, 2017; Polak et al., 2018; Povel, 1981; Ravignani et al., 2016; Repp et al., 2011, 2012; Schulze, 1989; Summers et al., 1986). Inversely, the two-category model best explaining adults’ neural responses did not significantly describe rhythm categorisation at birth. These differences suggest significant tuning of rhythm categories across the lifespan.

In our experiment, newborns’ data suggested the presence of an additional rhythm category that did not include the smallest integer ratios (i.e., not including 1:1 nor 2:1). This difference could be due to a process of learning and plasticity which may have tuned rhythm categorisation to socially relevant and statistically regular structures of the music-cultural environment (Polak et al., 2026). Specifically, in the current experiment, we tested Western Europe-enculturated adult participants (i.e., the most common cultural background given the location of the laboratory facilities), thus with expected experience mainly with Euro-American art music and global popular music. Notably, these music repertoires feature a prevalence of rhythms based on the smallest integer ratios (1:1 and 2:1; e.g., see Condit-Schultz, 2017; De Clercq, 2016; Temperley, 2010), in contrast with other music repertoires that prominently comprise also larger integer ratio rhythms (e.g., in Eastern Europe, South Asia, the Middle East and Africa; Brailoiu, 1951; Clayton, 2020; Cler, 1994; Goldberg, 2016; Johansson, 2017; Kvifte, 2007; Polak, 2010; Polak & London, 2014). A recent global survey of rhythm categorisation found corresponding categories in groups from these very regions, suggesting strong correlations between collective cultural practices and individual perceptual systems (Jacoby et al., 2024; Polak et al., 2026). It could thus be speculated that, in our case, the higher-integer-based rhythm category observed at birth may have been merged across the lifespan with the smaller-integer-based ones, reflecting a narrowing process driven by prevalent experience with the latter types of rhythm in the environment of the adult participants.

The narrowing of perceptual categorisation by the social relevance and statistics of the environment has been shown in several domains such as speech or vision. For instance, young infants can discriminate both native and non-native speech sounds, but their ability to discriminate among foreign speech sounds decreases already during the first year of life (Kuhl et al., 1992; Werker & Tees, 1984). In the visual domain, while 6-month-old infants can distinguish different human faces as well as different monkey faces, 9-month-old infants discriminate only faces of human individuals (Pascalis et al., 2002). Likewise, the progressive tuning of rhythm abilities according to the social prominence and statistical regularities of the musical environment has been shown in early childhood (for review, see Polak et al., 2026; Trainor & Hannon, 2013). For example, while musically untrained Western-enculturated adults are better at detecting changes in melodic patterns based on native than non-native scales, 6-months old infants performed equally well in both cases (Lynch et al., 1990). Moreover, at 6-to 7-month of age infants can detect structure violations in rhythmic patterns from familiar and unfamiliar musical systems (Hannon & Trehub, 2005a), in contrast with 12-month-old infants who, as adults, show a bias for culturally familiar rhythms in this violation detection (Hannon & Trehub, 2005b), a process interpreted as perceptual narrowing. Our results, by showing that the number of rhythm categories observed in newborns is narrowed down in adults, are thus compatible with the view that categories that are less or not used would progressively be lost over development (Maurer & Werker, 2014), a process that has been related to activity-dependent synaptic pruning (Faust et al., 2021; Kral, 2013).

However, it is important to note that the differences between newborns’ and adults’ neural responses in the present study were also contingent to the cultural familiarity of the adults tested here (i.e., Western Europe-enculturated participants). In other words, whether the narrowing observed between newborns and adults would hold with adult participants with a different music-cultural background (e.g., music-cultural environment with pervasive larger integer ratio rhythms) remains to be investigated. At the current stage of knowledge, we thus advocate for a more parsimonious view of plasticity whereby rhythm categories that are relevant to interaction with the environment would be selected and fine-tuned over development, without exclusively, or necessarily, entailing a process of selection by elimination or "narrowing".

### Exploring the role of prenatal experience in rhythm categorisation

In the present study, we tested newborns during their first to third post-natal day to minimise the effect of experience with the music-cultural environment. However, it could be argued that the rhythm categories measured at birth do not reflect neurobiological predispositions alone, as they are not necessarily independent from experience. Indeed, the human auditory system starts to respond to sounds as early as the last trimester of gestation (Daneshvarfard et al., 2020; Ghio et al., 2021; Mahmoudzadeh et al., 2016; Parncutt, 2016; Rotteveel et al., 1987; Saadatmehr et al., 2025), and the prenatal environment is rich in rhythmic structure from acoustic, vestibular, proprioceptive, and tactile stimulations. These include the fetal and maternal heartbeats, body movements, the mother’s respiration, speech, locomotion, and sensory inputs from external sources in the environments (James, 2010; Parga et al., 2018; Reid & Dunn, 2021; Richards et al., 1992; Voegtline et al., 2013). Prenatal rhythms have been suggested to play a critical role in shaping the processing of rhythm and music as further experienced over the lifespan (Parncutt, 2016; Parncutt & Chuckrow, 2019). In this context, we could speculate that the additional larger-integer ratio rhythm category observed in newborns as compared to adults could arise from prominent experience with similar rhythmic ratios in the intra-uterine environment. The heartbeat could be a candidate for such prenatal prominent rhythm, as the systole and diastole durations of the cardiac cycle typically correspond – depending on heart-rate – to such lager-integer ratios (e.g., systole-diastole ratios at rest often fall in the vicinity of 3:2; Bombardini et al., 2011; Hall et al., 2021).

Hence, our results could serve as a basis to study rhythm categorisation while reducing even more drastically the potential influence of prenatal experience, for example by testing pre-term newborns (Edalati et al., 2022, 2023, 2024; Saadatmehr et al., 2025) or fetuses (Corvilain et al., 2025). Complementarily, future research could compare neural categorisation of rhythm across different newborns groups with systematic differences in prenatal experience of rhythm, to understand whether and what environmental factors may contribute to shaping rhythm categorisation across different ages and cultures.

### Methodological advances and limitations of the current study

Task-free EEG recordings combined with the fRSA approach (Barbero, Lenc, et al. 2025) presents several advantages to reveal rhythm categorisation at birth. First, by moving from a unit-specific (here, amplitude of neural responses) to unit-free representational geometries (here, representational similarity matrices; Gennari et al., 2023; Grootswagers et al., 2022; Kriegeskorte, 2008; Kriegeskorte & Kievit, 2013; Robinson et al., 2025; Xie et al., 2020), fRSA enables direct quantification of the convergence of the results across age groups and measures (neural versus tapping responses in adults; Supplementary Material), irrespective of nonspecific differences in absolute amplitude of the responses across these groups and measures. Second, cross-age comparisons often require adapting the experimental paradigm in terms of task, especially in the case of testing preverbal infants, entailing the comparisons of measures that might be hard to directly compare (e.g., discrimination tasks in adults versus preferential looking time in infants). Our approach relying on task-free EEG recording of brain activity allows to overcome this limitation, and to control for confounds potentially stemming from decisional or motor biases. This is once again particularly critical for further comparing the responses across different age groups.

However, the approach used in the current study also presents some limitations, such as the lack of information on the impact of the newborns’ exact sleep state. Testing newborns during sleep is common practice in magneto/electro-encephalography (M/EEG) experiments, including those investigating rhythmic and auditory processes (Bianco et al., 2026; Edalati et al., 2022, 2023, 2024; Háden et al., 2015, 2024; Kushnerenko et al., 2007; Moser et al., 2020, p. 202; Ortiz Barajas et al., 2021; Panzani et al., 2023; Saadatmehr et al., 2025; Winkler et al., 2009). However, whether neural responses to rhythm differ when infants are in active state or not remains unknown (see for example Moser et al., 2020, for significantly different MEG responses to an auditory paradigm in infants up to 2 months of age in an active vs. sleep state). Moreover, the neural responses obtained from newborns were averaged to increase the signal-to-noise ratio of the responses by reducing the contribution of activity not temporally aligned across participants. While yielding a more stable representation, this procedure is at the expense of information about inter-individual variability (but see also Supplementary Results for analysis considering newborns’ individual data).

### Conclusions

Our findings indicate that human newborns, only a few days after birth, already exhibit neural representations consistent with rhythm categorisation. These results suggest the presence of neurobiological predispositions supporting the species-typical ability to categorise rhythm from the outset of life. These early processes likely form a foundation upon which experience later shapes rhythm processing through learning and plasticity, linking the biological roots of temporal processing to the diversity of music practices encountered worldwide.

## Materials and Methods

### Experimental model and study participant details

We tested two groups of participants: a group of 0- to 3-days old human newborns and a group of adults. Forty healthy human newborns participated in the study (21 females); two newborns were excluded from further analyses as one newborn was fed during the experiment and another newborn was born before 37 weeks of gestational age. Hence, the final sample included 38 healthy full-term newborns (20 females, age range at testing: 0-3 days, mean age ± SD at testing: 1.97 ± 0.43 days; gestational age range at birth: 37.29 – 41.57 weeks, mean gestational age ± SD at birth: 39.70 ± 1.06 weeks). All newborns had appropriate birth weight (mean weight ± SD = 3.36 ± 0.45 kg, range: 2.43 - 4.18 kg), head circumference (mean ± SD = 34.92 ± 1.39 cm, range: 32 – 37 cm – data available for all but two participants), and normal clinical neurological assessments. At least one of the parents were informed about the study and provided their written informed consent. The experimental procedure was approved by the ethics committee (Comité de Protection des Personnes Ouest I, France, ID-RCB: 2019-A01534-53).

The adult group comprised 19 participants (13 females, mean age ± SD = 25.57 ± 5.13 years, range: 19.29 – 38.26 years) with various levels of musical and dance training (musical training, mean ± SD: 4.05 ± 6.11 years, range: 0 - 18 years, 12 participants never had any musical training; dance training, mean ± SD: 0.11 ± 0.46 years, range: 0 - 2 years, 18 participants never had any dance training). Sample size was chosen in accordance with previous studies investigating rhythm perception using fRSA and/or frequency domain analyses (Barbero, Lenc, et al. 2025; Lenc et al., 2018; Nozaradan et al., 2011, 2012, 2018). As previous studies have shown an effect of music-cultural environment on rhythm perception (Hannon, Soley, et al., 2012; Hannon, Vanden Bosch der Nederlanden, et al., 2012; Hannon & Trehub, 2005; Jacoby et al., 2024; Jacoby & McDermott, 2017; Polak et al., 2018, 2026; Ullal-Gupta et al., 2014), adult participants were recruited in an effort to minimise variability in music enculturation. We tested adults who had had prominent experience with Western music and culture (participants were recruited in Brussels, Belgium), using country of origin and familiarity with music and cultures as a proxy for music-cultural experience. To participate in the study, adult participants had to self-report that they had lived, for at least the first 15 years of their lives, in one of the following countries: Belgium, France, United Kingdom, Netherlands, Luxembourg, Germany, Spain, Portugal, Italy or Switzerland (Guérin et al., 2024). Participants also had to self-report that they were not familiar with music and/or culture of countries other than those included in the aforementioned list. All participants reported to have normal hearing and no history of auditory or neurological disorders. The experimental procedure was approved by the local ethical committee at UCLouvain (project 2018-353). Adult participants provided their written informed consent to participate in the study and received financial compensation. The experimental procedures were conducted in accordance with the Declaration of Helsinki.

### Method details

#### Stimuli

The stimuli consisted of two-interval rhythmic patterns generated using Matlab R2022a (MathWorks). Such two-interval rhythmic patterns are formed by three successive acoustic events (here, an identical conga drum sample). The first temporal interval (IOI 1) is delimited by the first and second acoustic onset, while the second temporal interval (IOI 2) is delimited by the second and the third acoustic onset. If the rhythmic pattern is looped over time, the third acoustic event of one repetition also constitutes the first event of the subsequent repetition of the pattern (Fig. 1A). Such two-interval rhythms can also be described by the ratio between the duration of the two constituent temporal intervals. For instance, if the first interval is twice as long as the second one, we can refer to that pattern as a 2:1 ratio rhythm.

We generated 10 different two-interval rhythmic patterns conveyed by a repeated conga drum sample. All rhythmic patterns had a fixed total duration of 0.75 s, and the position of the second event was varied to obtain interval ratios over a temporal continuum between 1:1 to 2:1 across the 10 rhythmic patterns (Fig. 1, Supplementary Table 1). Each rhythmic pattern was individually looped over time to obtain 30-s long rhythmic sequences (i.e., 40 patterns repetition), each characterised by an interval ratio. The obtained rhythmic sequences (from the 10 distinct rhythmic patterns) constituted the 10 experimental conditions.

#### Experimental procedure in the newborn group

Newborn participants were presented with the 10 rhythmic sequences, each repeated four times in a pseudorandomised order (i.e., 40 trials). Trials were presented one after the other with a variable inter-trial interval between 4 to 6 seconds. The experiment could be interrupted at any moment in case of any sign of discomfort from the participants or their caregivers. Scalp electroencephalography (EEG) was recorded during sleep at the maternity ward of the Amiens University Hospital in France. We recorded EEG responses to all 40 trials in 36 participants. Due to technical issues during recording (missing triggers), we recorded 39 trials in one participant and 36 trials in another participant. Newborn participants were presented with the rhythmic sequences while lying in a cradle (N = 18) or in the caregiver’s arms (N = 19; caregivers were instructed not to move during the experiment), with this information missing for one of the tested newborns. The experiment was implemented in Matlab 2014b (MathWorks) using the Psychophysics Toolbox extensions (Brainard, 1997; Kleiner et al., 2007; Pelli, 1997). The stimulus sequences were presented via loudspeakers at a comfortable hearing level between 60-70 dB SPL. The experiment had an overall duration of approximately 23 minutes.

#### Experimental procedure in the adult group

The experimental procedure adopted for the adult group consisted of 4 listening blocks and 3 tapping blocks presented in alternation (see Supplementary Material for behavioural responses). In each block, the 10 rhythmic sequences were presented once in a randomised order.

During the listening blocks, participants were instructed to focus on the sequences, avoid any unnecessary movement and muscular tension, and fixate a cross displayed in front of them to minimise muscular and ocular artefacts in the EEG signal. To ensure focus to the rhythmic sequences, participants were instructed to detect transient intensity drops of the drum sounds within each sequence. Specifically, these transient intensity drops consisted in a 15% decrease in amplitude of the drum sounds over four consecutive repetitions of the rhythmic pattern within a sequence. In each sequence, there could be either one drop, two drops or none (number of occurrences over the 4 presentations across all listening blocks: 1, 1, or 2, respectively; instruction: “You will need to listen carefully to the audio sequences while fixating the fixation cross in front of you. Your task is to detect any occasional drop in volume in the sequences, which you will report at the end of each trial. In each sequence, there can be no drops, one drop, or two drops in volume”). Participants reported verbally the number of perceived intensity drops at the end of each sequence and received immediate feedback.

The experiment was implemented in Matlab R2016b (MathWorks) using the Psychophysics Toolbox extensions (Brainard, 1997; Kleiner et al., 2007; Pelli, 1997). The rhythmic sequences were presented binaurally through insert earphones (ER-2, Etymotic Research, Elk Grove Village, IL; air-conducted sound from the level of the participants’ clavicle to decrease magnetic interferences) connected to a Fireface UC audio interface (RME Audio, Haimhausen, Germany; sampling frequency = 44100 Hz) at a level of 73.2 dB SPL. Participants performed the experiment while comfortably seated in a chair. The experiment had an overall duration of approximately 70 minutes including the EEG and the behavioural sessions, and optional breaks between blocks.

#### EEG recordings

Brain activity of newborns was recorded using a 124-channels HydroCel Geodesic Sensor Net. The EEG signal was digitised at 1000 Hz with a Cz vertex electrode as a reference.

Brain activity of adults was recorded using a 64-channels BioSemi Active Two EEG system (https://www.biosemi.com/Products_ActiveTwo.htm) with two additional electrodes located on the left and right mastoids. Recording sites included standard 10-20 system locations (channel coordinates: https://www.biosemi.com/headcap.htm). Electrodes offset relative to the common mode sense (CMS) and driven leg (DRL) electrode loop was kept below ± 50 mV. An accelerometer was placed on the head of the participants to monitor whether participants complied with the instruction to avoid moving during the listening blocks. The signals from all channels and the accelerometer were digitised at 1024 Hz.

#### Quantification and statistical analysis

Neural data were analysed using Letswave 6, Letswave 7 (https://github.com/NOCIONS/letswave6, https://github.com/NOCIONS/letswave7) and custom-built scripts running in Matlab R2022a (Mathworks).

#### EEG data preprocessing in the newborn group

We discarded channels located at the external ring of the electrode net as they are more prone to contamination from movement and muscle artefacts, and they capture neural activity in response to auditory stimuli to a lesser extent as compared to central electrodes (Cirelli et al., 2016; Lenc et al., 2023).We applied a fourth order highpass Butterworth filter (cut-off at 0.5 Hz) and a fourth order lowpass Butterworth filter (cut-off at 30 Hz) to raw continuous EEG data, to remove slow drifts and higher frequencies irrelevant to the current study. We segmented the continuous data from 5 s before onset to 5 s after offset of the auditory stimulation. We then applied the Artifact Blocking algorithm (Mourad et al., 2007) to remove artefacts with amplitude exceeding ± 100 μV. The Artifact Blocking algorithm is often used in infant research to improve signal-to-noise ratio in continuous data (Calce et al., 2024; Cirelli et al., 2016; Edalati et al., 2024; Fujioka et al., 2011; Leleu et al., 2020; Lenc et al., 2023; Marie & Trainor, 2013, 2014; Rekow et al., 2021). Following visual inspection of the data (considering the data both before and after the implementation of the Artifact Blocking procedure), we linearly interpolated noisy electrodes with the three closest neighbouring channels (1 electrode in 3 newborns, 2 electrodes in 6 newborns, 3 electrodes in 1 newborns, 4 electrodes in 1 newborn, 5 electrodes in 2 newborns, 6 electrodes in 1 newborn, 7 electrodes in 2 newborns, 16 electrodes in 1 newborn, 21 electrodes in 1 newborn, and no electrode in the remaining 20 newborns), and we discarded segments containing excessive artefacts while retaining at least one segment per condition in each newborn (number of segments retained, mean ± standard deviation = 32.55 ± 5.96, range: 19 - 40).

After artefact rejection, we removed some additional electrodes located at the periphery of the net, keeping 98 out of the initial 124 electrodes (in line with Edalati et al., 2023). Data were re-segmented from 0 s to 30 s relative to the onset of the auditory stimulation (i.e., corresponding to trial duration). This second segmentation ensured that resulting epochs had a duration corresponding to an exact integer multiple of the rhythmic patterns’ duration, to avoid data overspill into frequency bins surrounding the frequencies of interest after applying the Fourier transform (Bach & Meigen, 1999). The data were then downsampled by a factor of 4 (i.e., to 250 Hz) and re-referenced to average mastoids to maximize the EEG responses to the acoustic stimuli (Luck, 2014; Nozaradan et al., 2018). Epochs were averaged in the time domain separately for each electrode, condition and participant. At this stage, the processing pipeline diverged for: (i) the identification of the frequencies of interest, (ii) the selection of participants based on the responsiveness to the auditory rhythmic stimulation, and (iii) the computations preceding the construction of the representational similarity matrix (RSM).

#### EEG frequency domain analysis in the newborn group

Following common practices in frequency domain designs (Barbero et al., 2021, 2025; Rekow et al., 2021; Retter, 2016; Volfart et al., 2021), we identified as frequencies of interest consecutive harmonics of the rhythmic pattern presentation frequency at which significant responses could be observed (i.e., at f = 1 / rhythmic pattern duration = 1 / 0.75 s = 1.33 Hz and harmonics: 2f = 2.66 Hz, 3f = 4 Hz, and so on). First, we averaged the time domain responses across participants, conditions, and a pool of fronto-central electrodes (channels: E4, E5, E6, E7, E11, E12, E13, E19, E20, E24, E28, E29, E30, E105, E106, E111, E112, E117, E118, E124, Fig. 2A). Electrode selection was based on the observation that responses to auditory rhythmic stimuli are best captured by fronto-central electrodes in infants (Cirelli et al., 2016; Edalati et al., 2022, 2023, 2024; Lenc et al., 2023; Saadatmehr et al., 2025) and in adults (Barbero, Lenc, et al. 2025; Lenc et al., 2018, 2020b; Nozaradan et al., 2012; Tierney & Kraus, 2015). We then submitted the data to a Fast Fourier Transform (FFT) to obtain a magnitude spectrum, and we computed the z-score at each frequency bin. Z-scores were calculated as the difference between the amplitude at the bin of interest (*x*) and the average amplitude of 8 surrounding bins (4 on each side, surrounding bins constituting a baseline: *bl*) excluding the immediately adjacent bins, divided by the standard deviation of the same selected bins 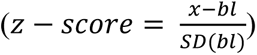. Given the frequency resolution of 0.0333 Hz (i.e., 1 / sequence duration), we considered surrounding bins up to ± 0.1667 Hz around each frequency of interest, excluding the immediately adjacent bins (see also: (Barbero et al., 2021; Retter & Rossion, 2016; Volfart et al., 2021). Responses at frequencies of interest were considered as significant if their z-score was higher than 3.09 (i.e., p < 0.001, one-tailed, signal > noise). An even number of consecutive significant frequencies of interest were considered for the following analysis to avoid an imbalance in the quantification of the response across conditions. Indeed, considering an odd number of harmonics would be at risk of under-representing responses to the 1:1 rhythm which is composed exclusively of even harmonics of the rhythmic pattern repetition frequency (i.e., the 1:1 rhythm corresponds to a frequency twice faster than the frequency of the pattern repetition).

We selected a subset of newborn participants for the fRSA analysis based on the responsiveness to auditory rhythmic stimulation regardless of condition. For each newborn, we averaged the time domain responses across fronto-central channels and conditions, and then we computed the magnitude spectra. We segmented each individual magnitude spectrum into chunks centred around the frequencies of interest identified from group-level data. Each chunk contained 11 bins: the frequency of interest and 5 bins on each side. The chunks were averaged, and a z-score was calculated as the difference between the amplitude at the bin of interest and the average amplitude of 8 surrounding bins (4 on each side excluding the immediately adjacent bins), divided by the standard deviation of the same surrounding bins. Newborn participants were retained for the fRSA analysis if their z-score was higher than 1.28 (i.e., p < 0.1, 1-tailed, signal > noise; Fig. S1). It is important to note that, as the z-scores for participant selection are calculated on individual data averaged across conditions, they are indicative of a response occurring at the rhythmic pattern presentation frequency and harmonics regardless of potential categorisation of rhythm.

To build the representational similarity matrix (RSM), we grand-averaged the data of the selected newborns to increase the signal-to-noise ratio (SNR) of the response (at the cost of losing individual variability) (Xie et al., 2022). The data was then averaged across the pool of fronto-central electrodes and submitted to a Fast Fourier transform (FFT) to obtain a magnitude spectrum for each condition. These responses were used to compute the representational similarity matrices.

#### EEG data preprocessing in the adult group

We applied a fourth order highpass Butterworth filter (cut-off at 0.1 Hz) and a fourth order lowpass Butterworth filter (cut-off at 64 Hz) to raw continuous EEG data to remove slow drifts and very high frequencies irrelevant to the current study. Continuous EEG data was then segmented from 5 s before onset to 5 s after offset of the auditory stimulation. Continuous EEG data was visually inspected for artefact rejection. At this stage, the data from two adults were discarded from further analysis. One adult was excluded due to the not compliance with the instruction of not moving during the listening blocks. Another adult was not included due to the presence of substantial artefacts on the EEG trace. Hence, the following analyses were performed on a sample of 17 adults. After visual inspection, we linearly interpolated noisy electrodes with the three closest neighbouring channels (1 electrode in 4 adults, 2 electrodes in 3 adults, 4 electrodes in 2 adults, no electrodes in the remaining 8 adults). Segments containing excessive artefacts were discarded (based on visual inspection of the data), retaining at least two segments per condition in each adult (number of discarded segments: mean ± standard deviation = 1.76 ± 3.54, range: 0 - 12).

To remove artefactual activity due to blinks and eye movements, we performed independent component analysis (ICA). ICA matrices were computed on data that underwent a slightly different preprocessing pipeline as compared to the one described above. We used a higher high-pass filter (cut-off 1 Hz, 4^th^ order Butterworth filter) to improve artefact classification accuracy (Chaumon et al., 2015). The data were segmented from 0 s to 30 s with respect to sequence onset (i.e., corresponding to trial duration) and downsampled of a factor of 4 (i.e., to 256 Hz) to increase ICA matrices computation speed. ICA matrices were obtained using the probabilistic ICA model with Laplace approximation. Artefactual independent components (ICs) due to blinks and eye movement were identified from visual inspection of the ICs time course and topographies, and then removed (1 IC removed in 16 participants, 2 ICs removed in 1 participant).

After artefact rejection data were re-segmented from 0 s to 30 s relative to the onset of the auditory stimulation (i.e., trial duration). The data were then downsampled by a factor of 4 (i.e., to 256 Hz), re-referenced to average mastoids, and averaged across trials (separately for each participant, condition, and electrode). The analytical pipeline then diverged for: (i) the identification of the frequencies of interest, and (ii) the preparation of the data for RSMs construction.

#### EEG frequency domain analysis in the adult group

To identify the frequencies of interest, the data was averaged across participants, conditions, and electrodes (excluding mastoids used for re-reference). We submitted the data to a Fast Fourier Transform (FFT) to obtain a magnitude spectrum, and we computed z-scores at each frequency bin as for the newborn data. We considered as frequencies of interest consecutive harmonics of the rhythmic pattern presentation frequency with a z-score higher than 3.09 (i.e., p < 0.001, one-tailed, signal > noise), and considering an even number of harmonics.

Individual preprocessed time domain data was averaged across a pool of fronto-central electrodes (channels: F1, Fz, F2, FC1, FCz, FC2, C1, Cz, C2, Fig. 2B) as responses to auditory rhythmic stimuli are best captured over fronto-central electrodes with scalp EEG (Barbero, Lenc, et al. 2025; Lenc et al., 2018, 2020b; Nozaradan et al., 2012; Tierney & Kraus, 2015).The data was then submitted to a Fast Fourier transform (FFT) yielding a magnitude spectrum for each participant and condition.

#### Auditory nerve modelling

We used a biologically plausible, well-established, forward model of the periphery of the auditory pathway (Bruce et al., 2018) to simulate the responses to the rhythmic sequences in a set of auditory nerve fibers. We implemented the simulation using the UR_EAR toolbox (version 2020b) running on MATLAB_R2022a (MathWorks) and the same parameters as in (Barbero, Lenc, et al. 2025). For each rhythmic sequence, we modelled responses from 128 cochlear channels with characteristic frequencies logarithmically space between 130 and 3000 Hz. The default cochlear tuning parameters matched data available from human participants (Shera et al., 2002). A biologically plausible distribution of high-, mid- and low-spontaneous rate fibers (respectively: 8, 12, and 31 – 51 in total) was simulated for each channel (Liberman, 1978). We then summed the resulting instantaneous firing rate across fibers and cochlear channels (Rajendran et al., 2020; Zuk et al., 2018) to obtain an estimate of the firing rate variation over the course of the sequences duration.

#### Representational similarity matrices

We analysed neural and behavioural responses using fRSA as part of the representational similarity analysis (RSA) framework (Kriegeskorte, 2008; Kriegeskorte & Kievit, 2013). RSA is a pattern information analysis consisting in the comparison of representational geometries that can be derived from stimuli descriptors, empirical data, and conceptual and computational models. Crucially, RSA moves from a unit-specific to a unit-free space, allowing to investigate whether brain responses might reflect stimulus features, higher-level categories, or aspects of overt behaviour.

To characterise the representational geometries of: (i) the neural responses, (ii) the auditory rhythmic stimuli, and (iii) the theoretical models of rhythm categorisation, we computed representational similarity matrices (RSMs) based on pairwise similarities across conditions. Hence, all RSMs were symmetrical around the diagonal (because of the non-directionality of the similarity across pairs of conditions) and had a value of 1 over the diagonal, reflecting maximal similarity of the response to one condition with itself. The procedure to build the representational geometries for the various descriptors is detailed here below.

For the grand-averaged newborn data, and for individual adult neural data, we used the amplitude values at the frequencies of interest to build a feature vector for each condition (Fig. 1C). Hence, the neural response in each condition was characterised by a feature vector containing *n* elements, *n* corresponding to the number of frequencies of interest. We then performed pairwise Pearson’s correlation across condition-specific feature vectors to build neural representational similarity matrices (Fig. 1D).

The representational geometry of the stimuli was described in three different ways by considering the temporal features (i.e., onset time) of the rhythmic patterns, the acoustic envelope of the stimuli, and the modelled auditory nerve responses.

The representational similarity matrix describing the temporal structure of the rhythmic stimuli was computed from inter-onset interval ratios. For each condition, the inter-onset interval ratio was calculated as the duration of the first time interval divided by the duration of the rhythmic pattern 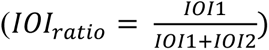. Similarity across conditions was quantified as the absolute difference between IOI ratios subtracted from a value of 1 (i.e., *similarity_m_*_−*n*_ = 1 −|*IOI ratio_m_* − *IOI ratio_n_*|; Fig. 1D). Hence, the stimulus RSM reflected the gradual dissimilarity across conditions moving along the 1:1 to 2:1 temporal continuum of the rhythmic sequences.

To build the acoustic envelope RSM, we extracted the envelope of the acoustic stimuli using the Hilbert transform and we submitted the envelopes’ time courses to a Fast Fourier transform. The RSM was obtained by performing pairwise Pearson’s correlations between magnitude responses at the frequencies of interest (as defined from EEG analysis).

The auditory nerve representational similarity matrix was computed from the modelled responses. For each condition, we submitted the simulated firing rate variation over time to a Fast Fourier Transform to obtain a magnitude spectrum. We then performed pairwise Pearson’s correlations across amplitude values at frequencies of interest (as defined from EEG analysis) to build a RSM (Fig. S5).

Theoretical models of rhythm categorisation were built to reflect the operational definition of categorisation: between-categories discrimination and within-categories generalisation (Rossion et al., 2015). In other words, categorical representational similarity matrices were designed to model the ability of the system to provide different responses to rhythms belonging to different categories and the same response to rhythms belonging to the same category. In the RSMs, similarity across conditions belonging to the same rhythm category was set to 1, while similarity across conditions belonging to different rhythm categories was set to 0. The number of categories within a model was not restricted a priori; however, each category had to include a minimum of two temporally contiguous conditions to comply with the within-category generalisation criterion. Having several theoretical models of rhythm categorisation presents the advantage of not having to a priori assume the number of rhythm categories and the locations of the categorical boundaries (i.e., the position where the switch from one rhythm category to another would occur). In turn, the flexibility to estimate the best-fitting theoretical model from each empirical RSM allows to quantify the consistency or variations of rhythm categories representations across individuals (adults), groups (infants and adults), and across neural and behavioural responses (adults, Supplementary Material). In total, considering all possible theoretical models given the operational definition of categorisation and the number of experimental conditions, we had 33 theoretical categorical models: 7 two-categories models, 15 three-categories models, 10 four-categories models and 1 five-categories model (Fig. S2). The shared variance across categorical models is described in Fig. S3.

#### Revealing rhythm categorisation from empirical data

To assess rhythm categorisation from empirical data, we performed partial correlations between the empirical RSMs and the theoretical models of rhythm categorisation while partialling out the stimulus RSM (i.e., the process was reiterated a number of times equal to the number of theoretical categorical models considered) (Barbero, Lenc, et al. 2025). The stimulus RSM was partialled out to account for any structure in the empirical RSMs that could have been merely driven by the rhythmic stimuli’s features. For each RSM and theoretical model, we performed Spearman’s partial correlations considering the lower triangular part of the matrices excluding the diagonal to avoid inflating correlation values (Mattioni et al., 2022). For each RSM, we derived a distribution of correlation coefficients across all theoretical models of rhythm categorisation, and the “winning” or “best” categorical model was defined as the one with the highest correlation coefficient (Fig. 1D).

The significance of (i) the grand-average newborn neural results and the (ii) individual neural results in adults was assessed with permutation testing (10000 iterations) to ensure that the correlation coefficient relative to the winning theoretical model was higher than what would be expected from chance. For each permutation iteration and RSM, we randomly shuffled the values of the lower triangular empirical RSM (excluding the diagonal, which was not included in the analysis) and computed the partial correlation between the shuffled empirical RSM and the theoretical categorical models while accounting for the stimulus RSM. The correlation value corresponding to the winning categorical model (relative to the shuffled data, i.e. the “winning” categorical model is redefined for each iteration) was stored, and these values were used to build a null distribution of correlation values for statistical testing. The p-value was calculated as the proportion of the observations in the null distribution that had a correlation value higher than the one obtained with the empirical data (Rezk et al., 2020). To correct for multiple comparisons (i.e., multiple theoretical models), p-values were adjusted using the Bonferroni correction for the 33 theoretical models.

We also investigated the significance of the adult results at the group level (permutation testing, 10000 iterations). For each iteration, we shuffled the individual RSMs (same shuffling across individual RSMs for a given iteration) and stored the correlation coefficient with the winning categorical model. The correlation coefficients were averaged across participants for each iteration, and the resulting values were used to build a null distribution of correlation values. Group-level p-value was calculated as the proportion of the observations in the null distribution with a value higher than the average correlation coefficient from the empirical data. We used Bonferroni correction to adjust the p-value for the 33 theoretical models.

#### Evaluation of shared representations between newborns and adults

To investigate the presence of shared neural representations of rhythm across newborns and adults, we compared the outputs of the fRSA analyses obtained from magnitude spectra across groups.

First, we correlated the grand-average newborn RSM with individual adult RSMs while partialling out the stimulus RSM to avoid that overlapping neural representational geometries across groups could be merely driven by low-level acoustic features. To this end, we performed Spearman’s partial correlations between the grand-average newborn RSM and each individual neural RSMs partialling out the stimulus RSM (considering the lower triangular parts of the matrices and excluding the diagonal). The significance of the results was assessed using permutation testing (10000 iterations) at the individual and at the group-level (following the same procedure described above). For each adult participant and iteration, we performed partial correlations between the shuffled grand-average newborn RSM and individual adult RSM while partialling out the stimulus RSM. From the obtained correlation coefficients, we built a null distribution for statistical testing, and a p-value was calculated as the proportion of observations in the null distribution that had a higher value than the observed correlation coefficient. Statistical significance at the group level was assessed following the same procedure. In this case, the null distribution was obtained by averaging the correlation coefficients across participants for each iteration.

Next, we compared the distribution of correlation coefficients across all possible theoretical models of rhythm categorisation for newborns and adults. For grand-average newborn data and for individual adult participant, we considered the vectors consisting of the correlation coefficients with each theoretical model of rhythm categorisation. The vectors were Fisher transformed and then correlated across groups: the grand-average newborn vector was Pearson’s correlated with the vectors of individual adult participants. The resulting correlation values were then Fisher transformed, and tested against 0 with a one-sample t test.

#### Hierarchical clustering

We implemented multidimensional scaling and hierarchical clustering as a qualitative analysis to explore the representational structure of rhythm categories in newborns’ and adults’ neural responses (Mattioni et al., 2020). For newborns’ and adults’ responses, we first computed representational dissimilarity matrices (RDMs) by subtracting their similarity matrices from a value of 1 (i.e., RDM = 1 – RSM, operation computed for each pixel). Hence, we got dissimilarity matrices with a value of 0 across the diagonal, and symmetrical values around the diagonal. The newborns’ RDM was computed from the RSM computed from grand-averaged data, considering magnitude responses at the frequencies of interest (same as for the fRSA analysis). The adults’ RDM was computed by first averaging the individual RSMs (based on the magnitudes of responses at the frequencies of interest), and then subtracting the resulting RSM from a value of 1. For each group, a hierarchical cluster tree was computed using the Matlab function ‘linkage’ (method: average), and clusters were defined with the Matlab function ‘cluster’, indicating a maximum of either 2, 3, 4, or 5 clusters (i.e., hierarchical clustering was performed four times for each group). We used multidimensional scaling with a dimensionality of 2 to visualise results.

## Data and materials availability

Analysis code to reproduce the results of this manuscript and the adult group data will be made publicly available upon publication.

## Author contributions

Conceptualization: F.M.B., T.L., S.N.

Methodology: F.M.B., T.L., S.N., S.M.

Software: F.M.B., T.L., A.G.

Validation: F.M.B., T.L., S.N. Formal analysis: F.M.B.

Investigation: F.M.B., A.G.

Resources: F.M.B., T.L., S.N., S.M., A.F.

Data curation: F.M.B.

Writing - original draft: F.M.B., S.N.

Writing - review and editing: F.M.B., T.L., A.G., N.J., R.P., M.V., A.F., S.M., S.N.

Visualization: F.M.B.

Supervision: S.N.

Project administration: S.N., S.M.

Funding acquisition: S.N., S.M.

