## Supplemental Material for "Neural categorisation of rhythm in human newborns"

### Neural categorisation of rhythm in human newborns
### Supplementary Material

#### Supplementary Results

##### Magnitude responses best capture rhythm categorisation from newborns' neural responses

Newborns' neural responses obtained from the magnitude spectra revealed significant categorisation in response to rhythmic patterns. Next, we tested if comparable categorisation could be obtained from the complex spectra and time domain representations of the neural responses, in line with prior work in adults (Barbero, Lenc, et al. 2025). Using complex Fourier coefficients or time domain data allows to include not only information about what the response looks like (i.e., the shape of the response to the repeated rhythmic pattern, as captured with magnitude spectra) but also information on when this response starts relative to the onset of the stimulus sequence (i.e., time-locking to a common temporal reference across conditions or not). Applying the same RSA analysis, the results did not reveal significant categorisation (complex spectra RSM: permutation test, $\rho = 0.4515$, $p_{Bonf} = 1$; time domain RSM: permutation test, $\rho = 0.3932$, $p_{Bonf} = 1$, Fig. S4). Nevertheless, the representational similarity matrices obtained from complex spectra and from time domain responses significantly correlated with the RSM obtained from magnitude spectra (partial correlations of RSMs controlling for the stimulus temporal information; magnitude versus complex spectra and time domain responses, respectively: permutation test: $\rho = 0.4773$, $p = 0.0006$, and $\rho = 0.4352$, $p = 0.0016$). These null results, in contrast with the positive results obtained with the magnitude spectra, thus suggest that the neural responses to the rhythms, while showing a categorical structure across conditions in its shape, do not show a consistent categorical structure in the time-locking to a temporal reference across conditions, thus yielding disruption of the categorical structure when this information is included in the analysis. Whether this higher variability in time-locking of the responses across conditions results from lower signal-to-noise ratio in newborns' as compared to adults' data, or results from weaker association with an experience-driven prior of specific alignment of the categorical responses to rhythm remains an open question.

##### Neural categorisation of rhythm from group-level average and individual newborns' data

In addition to the results considering the representational similarity matrix (RSM) built from the neural responses averaged across 19 newborns, we performed group-level statistics considering individual newborns' RSMs, as conducted for the adults' response. We computed individual RSMs considering the magnitude of the response at the frequencies of interest (i.e., first 6 harmonics of the rhythmic pattern) distributed over fronto-central electrodes. We performed partial correlations between the RSMs and theoretical models of rhythm categorisation while controlling for the temporal information carried by the stimuli. Neural responses significantly correlated with theoretical models of rhythm categorisation at the group level (permutation test, $\rho = 0.3857$, $p_{Bonf} = 0.0033$, results significant at the individual level in

3 out of 19 newborns, $ps_{Bonf} < 0.05$), in line with results obtained from the data averaged across newborns.

#### Neural categorisation of rhythm in adults considering all harmonics

To complement the analysis of adult neural responses including the first 6 consecutive harmonics of the rhythmic pattern presentation frequency, we performed fRSA considering magnitude spectra responses at all harmonics up to 61.33 Hz (i.e., 46 harmonics, including all harmonics not attenuated by the 64 Hz low-pass filter). Neural RSMs significantly correlated with theoretical models of rhythm categorisation while partialling out the temporal stimulus RSM (permutation test, $\rho = 0.5538$, $p_{Bonf} = 0.0033$; results significant at the individual level in 10 out of 17 adults, $ps_{Bonf} < 0.05$). Results built from all harmonics were highly consistent with results obtained considering responses up to 8 Hz (i.e., 46 or 6 harmonics). In fact, there was a significant correlation between individual RSMs after partialling out stimuli information (permutation test, $\rho = 0.8727$, $p = 0.0001$; results significant in all participants, $ps < 0.05$), and a significant correlation between the distribution of correlation coefficients across all theoretical models of rhythm categorisation (mean $\rho$ across participants = 0.9285, one-sample t-test against zero, $p < 0.0001$). Moreover, there was no statistically significant difference between the correlation coefficients of the RSMs with the best-fitting categorical models when considering 6 versus 46 harmonics (paired t-test of Fisher-transformed correlation coefficients, $p = 0.7069$, Bayes factor $BF_{10} = 0.2658$, indicating moderate evidence for the null). Overall, these results suggest that considering the first 6 consecutive harmonics efficiently captured rhythm categorisation in adults' neural responses.

#### Neural categorisation of rhythm in adults obtained from complex spectra and time domain responses

Neural categorisation of rhythm was assessed across different analyses of the adults' neural responses. We observed evidence for neural categorisation of rhythm when we applied the same analytical pipeline to (i) the complex-valued EEG spectra (i.e., including the real and imaginary Fourier coefficients; permutation test, $\rho = 0.5465$, $p_{Bonf} = 0.0066$; results significant at the individual level in 9 out of 17 participants, $ps_{Bonf} < 0.05$), and to (ii) the time domain data (low-pass filtered at 10 Hz; permutation test, $\rho = 0.5502$, $p_{Bonf} = 0.0066$; results significant at the individual level in 10 out of 17 participants, $ps_{Bonf} < 0.05$; Fig. S4; stimulus RSM derived from the stimuli temporal structure, as for the main analysis).

Results obtained across different EEG measures (magnitude spectra, complex spectra, and time domain data) significantly correlated within participants. Convergence of the fRSA outputs across measures was assessed in different ways: (i) by correlating individual neural RSMs built from different measures while partialling out the stimulus RSM, (ii) by correlating the distribution of correlation coefficients obtained across all categorical models, and (iii) by comparing the correlation coefficients between the RSMs and the best-fitting categorical models.

The RSMs built from magnitude and complex spectra significantly correlated beyond the shared information driven by the stimulus temporal features (permutation test, $\rho = 0.5479$, $p = 0.0001$; results significant at the individual level in 15 out of 17 participants, $ps < 0.05$).

We observed a significant correlation between the distribution of correlation coefficient across the two measures (mean ρ across participants = 0.6722, one-sample t test against zero, p = 0.000005). The correlation coefficients of the RSMs with the best-fitting categorical models did not differ between magnitude and complex spectra responses (paired t-test of Fisher-transformed correlation coefficients, p = 0.6151, $BF_{10}$ = 0.2798, indicating moderate evidence for the null).

We identified significant correlations between fRSA results obtained from magnitude spectra and time domain data both at the level of neural RSMs (permutation test, ρ = 0.5689, p = 0.0001; results significant at the individual level in 16 out of 17 participants, ps < 0.05), and for the distribution of correlation coefficients across measures (mean ρ across participants = 0.6999, one-sample t test against zero, p = 0.000002). There was no significant difference between the correlation coefficients obtained between the RSMs and the best-fitting categorical models between magnitude spectra and time domain responses (paired t-test of Fisher-transformed correlation coefficients, p = 0.7505, $BF_{10}$ = 0.2609, moderate evidence for the null).

Altogether, our results suggest that, in the presence of EEG responses with higher signal-to-noise ratio (as for the adult data here), different measures appear to carry convergent information on the internal representation of rhythmic patterns, as the measures are less impacted by noise.

#### Convergence across behavioural and neural categorisation of rhythm in adults

In order to have a behavioural measure of rhythm categorisation, adult participants were instructed to tap their finger in synchrony with the rhythmic sequences in blocks separate from the EEG recordings. Continuous tapping force was recorded with a device, hereafter referred to as the tapping box, and the resulting signal was analysed using fRSA. Individual RSMs were built from: (i) inter-tap interval (ITI) ratios extracted from the continuous tapping force and (ii) the raw continuous tapping force [see also (Barbero, Lenc, et al., 2025) for other convergent analyses derived from the tapping signal].

We extracted tap onset times from continuous tapping data (Fig. S5), and we calculated the inter-tap interval (ITI) ratios. For each repetition of the rhythmic pattern, the ITI ratio was obtained by dividing the first ITI over the sum of the first and second ITI as in previous studies (Barbero, Lenc, et al., 2025; Polak et al., 2018). ITI ratios were then averaged across rhythmic patterns repetitions separately for each condition and participant (mean number of ITI ratios per condition = 99.8588, range = 47 – 114). For each participant, we built a RSM considering the absolute difference across the averaged ITI ratios produced in each condition. We then performed partial correlations between the RSM and theoretical models of rhythm categorisation while accounting for the temporal information carried by the stimuli. Our results revealed significant evidence of rhythm categorisation during sensorimotor reproduction of the rhythmic stimuli (permutation test, ρ = 0.7669, $p_{Bonf}$ = 0.0033; results significant at the individual level in all 17 adult participants, $ps_{Bonf}$ < 0.0033; Fig. S5C), in line with previous results obtained using various behavioural tasks [e.g., identification, discrimination, and reproduction (Clarke, 1987; Desain & Honing, 2003; Jacoby & McDermott, 2017; Nave et al.,

2024; Polak et al., 2018; Povel, 1981; Ravignani et al., 2016; Repp et al., 2011, 2012; Schulze, 1989; Summers et al., 1986)].

Next, we analysed the continuous tapping force. As for the neural data, we first identified the number of significant harmonics of the rhythmic pattern. We averaged the time domain continuous tapping force across participants and conditions. We then submitted the data to a Fast Fourier transform to obtain a magnitude spectrum, and we calculated the z-score value at each frequency bin. The magnitude of the response was significantly above the local noise baseline for all consecutive harmonics up to 6.67 Hz (i.e., the fifth harmonic; $z > 3.09$, corresponding to $p < 0.001$, one-tailed, signal > noise). Hence, further analyses were performed considering the first 6 harmonics as frequencies of interest (as responses to the 1:1 rhythm are expected to be observed only at even harmonics of the rhythmic pattern presentation frequency, considering an odd number of harmonics would bias the results).

We built individual RSMs from the continuous tapping force by performing pairwise correlations between the condition-specific magnitude of the responses at the frequencies of interest. We then performed partial correlations between the tap-force RSMs and theoretical models of rhythm categorisation while partialling out the temporal information carried by the stimuli, observing evidence in favour of rhythm categorisation (permutation test, $\rho = 0.7483$, $p_{Bonf} = 0.0033$; results significant at the individual level in all participants, $ps_{Bonf} < 0.05$; Fig. S5D). We observed evidence for rhythm categorisation from the continuous tapping force also when fRSA was applied to: (i) responses considering the complex FFT spectra (permutation test, $\rho = 0.7376$, $p_{Bonf} = 0.0033$; results significant at the individual level in all participants, $ps_{Bonf} < 0.05$), and (ii) the continuous tapping force in the time domain (permutation test, $\rho = 0.7450$, $p_{Bonf} = 0.0033$; results significant at the individual level in all participants, $ps_{Bonf} < 0.05$; Fig. S3).

Crucially, the results obtained from the continuous tapping force converged with the results obtained from ITI ratios. The RSMs obtained from ITI ratios and from the magnitude response of the continuous tapping force significantly correlated at the individual level after accounting for the temporal information carried by the stimuli (permutation test, $\rho = 0.8153$, $p = 0.0001$; results significant at the individual level in all participants, $ps < 0.05$). Our analyses also highlighted a significant correlation between the distribution of correlation coefficients across all theoretical models obtained from the two measures (mean $\rho$ across participants = 0.9313, one-sample t test against zero, $p < 0.0001$). There was no significant difference between the correlation coefficients obtained with the RSMs derived from magnitude spectra and ITI ratios (paired t-test of Fisher-transformed correlation coefficients, $p = 0.2031$, $BF_{10} = 0.5264$, indicating anecdotal evidence for the null). This result underscores the ability of fRSA to reveal rhythm categorisation from continuous data, a crucial advantage when identifying clear temporal markers in continuous signal is not straightforward, as for the neural responses to rhythm recorded in the current study (Barbero, Lenc, et al., 2025).

Lastly, we investigated the overlap between the neural and behavioural representational geometries recorded in adults. To do so, we correlated the fRSA outputs obtained from the magnitude spectra of the responses for brain and behaviour. The individual neural and behavioural RSMs (Fig. S6) were significantly correlated after removing the common information that could have been merely driven by stimuli onset time (permutation test, $\rho$ =

0.3383, p = 0.00009; results significant in 12 out of 17 participants, ps < 0.05; Fig. S5E). We also observed a convergence of the results when we correlated the distribution of correlation coefficients obtained for each participant for all theoretical model of rhythm categorisation (mean ρ across participants = 0.5048, one-sample t test against zero, p = 0.0002; Fig. S5F, G).

As when comparing neural responses in newborns and adults, we also investigated whether the categorical model best explaining neural data could efficiently capture rhythm categorisation from tapping responses, and vice versa. The categorical models that best explained adult neural and behavioural data were two-categories theoretical models characterised by a bigger rhythm category for uneven rhythms (neural responses: model 2, winning model in 6 out of 17 adults; behavioural responses: model 3, winning model in 8 out of 17 adults). When we performed partial correlations between the neural RSMs and the categorical model best explaining behavioural responses while accounting for the stimuli temporal information, we observed significant results (permutation test, ρ = 0.2426, p = 0.0179, results significant in 7 out of 17 adults, ps < 0.05). Results were also significant when performing partial correlations between tapping RSMs and the categorical model best explaining neural responses (permutation test, ρ = 0.5336, p = 0.0001, results significant in 13 out of 17 adults, ps < 0.05). The convergence between neural and behavioural responses to the same rhythms in adults was also observed using hierarchical clustering and multidimensional scaling. As for the neural responses, when hierarchical clustering was stopped at 2 clusters, we observed a correspondence with the theoretical model best fitting the behavioural responses (Fig. S7). When progressively increasing the number of possible clusters to 3, 4, and 5 to fit the data, clusters consisting of only one rhythmic condition appeared, suggesting that increasing the number of clusters might result in over partitioning (i.e., overfitting). Lastly, we observed that within-category rhythmic conditions clustered more closely together in tapping than in neural responses. This is not surprising, as recorded neural responses consists of a mixture of rhythm categorisation and tracking of the stimuli temporal features (note that in fRSA the contribution of the temporal tracking of stimuli features is controlled by using partial correlations). Together, these results highlight the consistency between the neural representation of the two-interval rhythms recorded in the absence of any motor- or timing-related task with the behavioural representations of the same rhythms measured during a sensorimotor synchronisation task [in line with results discussed in (Barbero, Lenc, et al., 2025)].

### Supplementary Methods

#### Experimental procedure – behavioural adult data

The experimental procedure consisted of 4 listening blocks and 3 tapping blocks presented in alternation. During each tapping block, the 10 rhythmic sequences from the morphed 1:1 – 2:1 temporal continuum (see Supplementary Table 1) were presented once in a randomised order. Participants were instructed to tap with the index finger of their preferred hand in synchrony with the auditory stimulation. Participants had to tap on a device (tapping box) recording both the continuous tap force signal and the tapping onsets. Participants were required not to tap before the beginning of the sequences, and not to wait too long to start tapping after sequences onset. For further details on experiment implementation and sound delivery please refer to: Materials and Methods – Experimental procedure in the adult group.

#### Behavioural recordings

Adult participants' tapping onsets and continuous tapping force were recorded using a custom-made device (tapping box) connected to the BioSemi Active Two EEG system's Analog Input Box (https://www.biosemi.com/aib.htm). Recorded tapping onsets, digitised as triggers, corresponded to the moments in time in which the finger got in contact with the conductive surface of the tapping box, hence closing an electrical circuit. The tapping force was recorded as a continuous signal using a six-axis force sensor (FT48224, ATI Industrial Automation, NC). The latency and jitter of the tapping signals were below 1 ms, as measured with an oscilloscope. Tapping signals (onsets' triggers and continuous force from the six sensors) were digitised with a sampling frequency of 1024 Hz. Additionally, we also recorded a copy of the auditory signal through the BioSemi Analog Input Box, which was digitised at 1024 Hz.

#### Quantification and statistical analysis

Behavioural data were analysed using Letswave 6, Letswave 7 (https://github.com/NOCIONS/letswave6, https://github.com/NOCIONS/letswave7) and custom-built scripts running on Matlab R2022a (Mathworks), following the analytical pipeline described in (Barbero, Lenc, et al., 2025).

##### *EEG time domain analysis in the newborn group*

The time domain analysis was performed on the subset of newborns selected based on the frequency domain analysis of neural responses (see Materials and Methods). We applied to the preprocessed data a fourth order Butterworth lowpass filter with a 10 Hz cut-off frequency. This cut-off frequency was used to match the frequency range corresponding to the neural responses to the stimuli, i.e., the range corresponding to the harmonics showing amplitude identified as being significantly above local noise in the magnitude spectrum of the responses. The filtered data was then segmented from 0 s to 30 s with respect to the onset of the auditory sequences (i.e., trial duration) and re-referenced to the averaged mastoids signal. At this stage, only the data from a pool of 20 fronto-central electrodes were retained for further analysis (Fig. 2A). The data was re-segmented to obtain epochs of the duration of the rhythmic patterns (i.e., 0.75 s), and downsampled by a factor of 10 to reduce dimensionality of the data (resulting sampling frequency = 100 Hz). For each participant, epoch, and electrode, the amplitude of the signal was demeaned by subtracting at each time point the average amplitude over the epoch to correct for possible offsets.

At this point, we implemented an automatic artefact rejection procedure to discard epochs containing excessive artefacts. Epochs where the range of the amplitude of the response exceeded a threshold of 100 μV in at least 25 % of the channels (i.e., at least 5 out of the 20 fronto-central electrodes of interest) were discarded from further analyses. Then, for each newborn the number of epochs was equalised across conditions to minimise differences in SNR across experimental conditions. The data was then averaged across participants (as for the frequency domain analysis, see Materials and Methods).

*EEG time domain analysis in the adult group*

After artefact rejection, we applied a fourth order Butterworth lowpass filter (10 Hz cut-off, based on the results from the frequency domain analysis). The data was segmented from 0 s to 30 s with respect to the onset of the auditory sequences (i.e., trial duration) and re-referenced to the averaged mastoid signal. Next, the data was re-segmented to obtain 0.75 s-long epochs (i.e., rhythmic pattern duration) and downsampled to a 64 Hz sampling rate. To correct for possible offsets, the data of each epoch was demeaned by subtracting at each time point the epoch's average amplitude separately for each epoch, electrode, and participant. The resulting epochs were then averaged separately for each participant and condition.

*Behavioural data preprocessing*

The continuous tapping force recorded from the six sensors of the tapping box was segmented from -1 s to 30 s relative to the onset of the auditory sequences. To correct for potential offsets present in the recording, the signal from each sensor was baseline corrected by subtracting at each time point the averaged signal over 1 s before trial onset. We then used the calibration matrix of the tapping box to compute the tapping force orthogonal to the surface of the box from the signal recorded from the six sensors. The obtained signal was then re-segmented to discard time points preceding the onset of the auditory stimulation and downsampled by a factor of 4 (resulting sampling frequency = 256 Hz).

*Tap onsets analysis*

Tap onset times were extracted from the continuous tapping force based on signal amplitude (i.e., surpassing a certain threshold) and minimum inter-tap interval (i.e., minimum time occurring between two events to be considered as valid tap onsets; Fig. 3A). Assigned tap onsets were visually inspected for validation; parameters could be adjusted on a subject-by-subject basis to optimise tap onset extraction.

Inter-tap interval ratios were calculated from tap onset times. A tap onset time was considered as valid for inter-tap interval ratio calculation if the tap was performed within a 0.124 s window centred around the onset time of a stimulus. If this condition was not met, a tap onset was discarded from further analysis. For each rhythmic pattern cycle, three valid tap onsets were needed for inter-tap interval ratio calculation: the taps corresponding to the first and second event, and the tap corresponding to the first event of the following rhythmic pattern repetition (to estimate second inter-tap interval duration). For each participant, condition, trial, and rhythmic pattern repetition, we calculated inter-tap interval ratio dividing the first inter-tap interval duration (ITI1) by the summed duration of the first and second (ITI2) inter-tap interval ($ITI_{ratio} = \frac{ITI1}{ITI1+ITI2}$; Fig. 3A). Inter-tap interval ratios were then summed across rhythmic pattern repetitions separately for conditions and participants (Barbero, Lenc, et al., 2025; Jacoby & McDermott, 2017; Polak et al., 2018).

*Behavioural data frequency domain analysis*

To determine the range of frequencies of interest, we followed the same procedure as for the neural data. We first averaged the time domain responses across participants and conditions. The data was then submitted to a Fast Fourier transform to obtain a magnitude spectrum. Then, we calculated z-scores at each frequency bin, considering as baseline responses at ± 0.1667 Hz, excluding the immediately adjacent bins (i.e., neighbouring bins 2-5 from both side; see also the procedure described in Materials and Methods - EEG frequency domain analysis). Consecutive harmonics with a z-score higher than 3.09 (i.e., $p < 0.001$, one-tailed, signal >noise) were considered significant and included in the fRSA analysis. As for the neural responses, we considered an even number of harmonics, to account for the 1:1 rhythmic pattern whose frequency content projects only onto even harmonics (each interval has a duration of 0.375 s, $f_{1:1} = 1 / 0.375$ s = 2.66 Hz = 2f).

To compute representational similarity matrices, the continuous tapping force was averaged in the time domain separately for each participant and condition.

##### *Behavioural data time domain analysis*

Informed by the frequency domain analysis, we applied a fourth order Butterworth lowpass filter with a 10 Hz cut-off to the continuous tapping force. We then re-segmented the data to obtain epochs with a length equivalent to the rhythmic pattern duration (0.75 s) and downsampled the data to a 64 Hz sampling rate to reduce dimensionality. We demeaned the data by subtracting at each time point the average amplitude of the signal in a given epoch to correct for possible offsets. The epochs were then averaged separately for each participant and condition.

##### *Representational similarity matrices*

Neural RSMs were built from magnitude spectra, complex spectra and time domain data (in all cases, from data averaged across fronto-central channels; see also Materials and Methods).

For the magnitude spectra RSMs, time domain responses were submitted to a Fast Fourier transform, and condition-specific features were built by considering the magnitude of the response at the frequencies of interest (feature vector dimension: $n$, with $n$ being the number of frequencies of interest). For the complex spectra RSMs, we calculated complex Fourier spectra and condition-specific feature vectors were built by concatenating the real and imaginary Fourier coefficients at the frequencies of interest [feature vector dimension: 2 (real and imaginary Fourier coefficient) x $n$ frequencies of interest]. For the time domain data, condition-specific feature vectors were obtained by concatenating the amplitude value at each time point over the course of the repetition pattern (feature vector dimension: $f_s \; x \; t$, with $f_s$ being the sampling frequency of the time domain data, and $t$ corresponding to the rhythmic pattern duration). For all measures, we performed pairwise Pearson's correlations across condition-specific feature vectors to obtain RSMs.

The representational similarity matrices (RSMs) of the behavioural responses were also based on pairwise similarities across conditions. Individual behavioural RSMs were built from magnitude spectra, complex spectra, and time domain data following the same procedure followed for the neural responses. Additionally, we also computed behavioural RSMs from ITI ratios. For each participant, the pairwise similarity across conditions was calculated by subtracting the absolute difference across produced ITI ratios from a value of 1 (i.e., $similarity_{m-n} = 1 - |ITI\ ratio_m - ITI\ ratio_n|$).

##### *Representational similarity analysis*

To assess rhythm categorisation from neural and behavioural data, we performed partial correlations between the empirical RSMs and the theoretical models of rhythm categorisation while partialling out the stimulus RSM. Significance of the results was assessed using permutation testing following the same procedure as outlined in Materials and Methods.

##### *Evaluation of shared representations between measures*

We compared the RSMs derived from magnitude spectra, complex spectra, and time domain representation of the newborns' neural responses using partial correlations while partialling out the stimulus RSM. Statistical significance was calculated using permutation testing (10000 iterations).

We also investigated the consistency of the responses across measures (i.e., neural and behavioural responses) and data descriptors (i.e., RSMs obtained from magnitude-, complex-spectra, and time domain for neural responses; RSMs obtained from magnitude spectra and ITI ratios for behavioural responses) of adult responses. Following the same rationale and procedures of the across groups comparisons, consistency across measures and data descriptors were investigated by: (i) comparing the representational similarity matrices, and (ii) comparing the distributions of correlation coefficients across different theoretical models of rhythm categorisation. Here, the RSMs and the vectors containing the correlation coefficients across theoretical models were correlated on a participant-by-participant basis. To investigate the similarity between RSMs, we performed partial correlations while partialling out the stimulus RSM; statistical significance was calculated using permutation testing (10000 iterations). Moreover, correlation coefficients across all possible models were correlated within-participants for the comparisons across measures. Statistical significance was assessed with one-sample t-tests against 0 (after Fisher-transforming the correlation coefficients; see also Materials and Methods). Lastly, we compared the correlation coefficients with the best-fitting model across individual neural measures in adults to test how prominent the observed categorical structure was between two kinds of responses. To do so, the individual Spearman's correlation coefficients with the best-fitting model were Fisher-transformed and compared across measures using a paired t-test. Bayes factors were calculated to quantify the evidence in favour of the alternative hypothesis over the null hypothesis ($BF_{10}$) using the bayesFactor package for MATLAB (https://github.com/klabhub/bayesFactor).

Additionally, hierarchical clustering of behavioural responses was implemented as for the neural responses (see Materials and Methods).

#### Supplementary Table

| Condition | IOI 1 [s] | IOI 2 [s] | IOI 1 ratio | IOI 2 ratio |
|---|---|---|---|---|
| **1 (1:1)** | 0.3750 | 0.3750 | 0.5000 | 0.5000 |
| **2** | 0.3889 | 0.3611 | 0.5185 | 0.4815 |
| **3** | 0.4028 | 0.3472 | 0.5370 | 0.4630 |
| **4** | 0.4167 | 0.3333 | 0.5556 | 0.4444 |
| **5** | 0.4306 | 0.3194 | 0.5741 | 0.4259 |
| **6** | 0.4444 | 0.3056 | 0.5926 | 0.4074 |
| **7** | 0.4583 | 0.2917 | 0.6111 | 0.3889 |
| **8** | 0.4722 | 0.2778 | 0.6296 | 0.3704 |
| **9** | 0.4861 | 0.2639 | 0.6481 | 0.3519 |
| **10 (2:1)** | 0.5000 | 0.2500 | 0.6667 | 0.3333 |

**Table S1. Inter-onset intervals of the stimuli across conditions.** Inter-onset intervals between the first and the second sound (IOI 1), and between the second sound and the first sound of the following repetition of the rhythmic pattern (IOI 2). The inter-onset intervals are also described in terms of the ratio between each IOI and the total duration of the rhythmic patterns. Values are shown separately for the 10 conditions, which were linearly covering the 1:1 - 2:1 temporal continuum.

#### Supplementary Figures

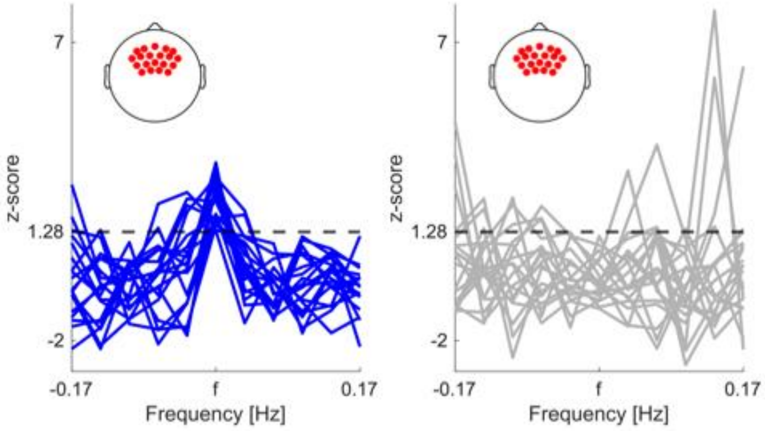


**Figure S 1. General auditory response to the rhythmic stimuli in individual newborn participants.** Z-score of the auditory response (f) and surrounding bins for newborn participants retained (left, in blue) and discarded (right, in grey) for further analyses. Z-scores were calculated from responses averaged across frequencies of interest (first 6 harmonics of the pattern repetition rate), fronto-central channels (highlighted in the insets), and conditions. Newborn participants were selected for further analyses if their z-score was higher than 1.28 (i.e., $p < 0.1$, 1-tailed, signal > noise; dashed line).

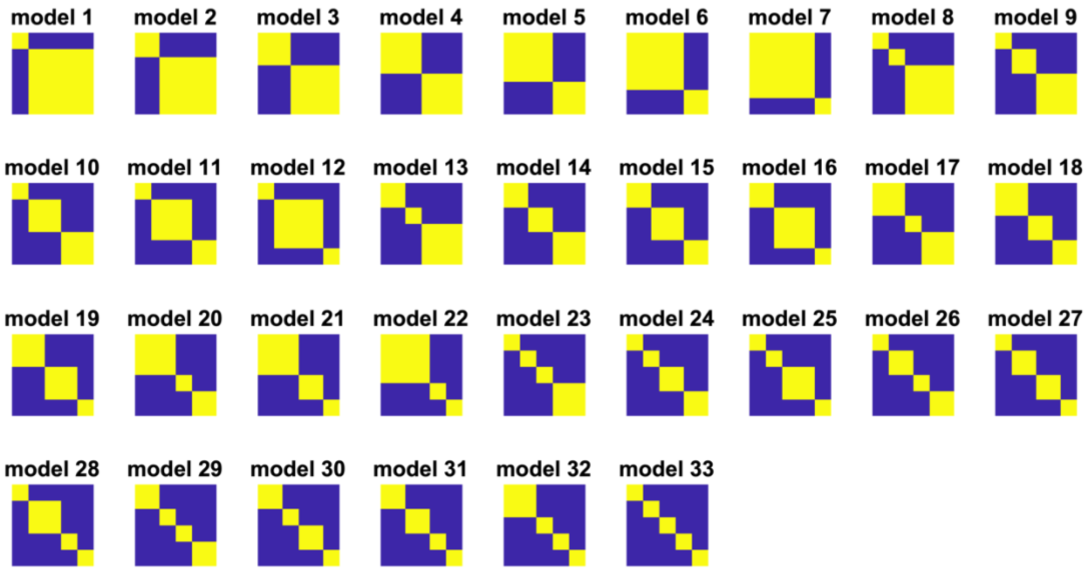


**Figure S 2. Theoretical models of rhythm categorisation.** The theoretical models consisted of all possible categorical models given the operational definition of categorisation (within-category similarity and between-categories dissimilarity) and the number of experimental conditions (here, 10 conditions). Therefore, each model consisted of at least two categories and each category consisted of at least two neighbouring conditions, for a total of 33 categorical models.

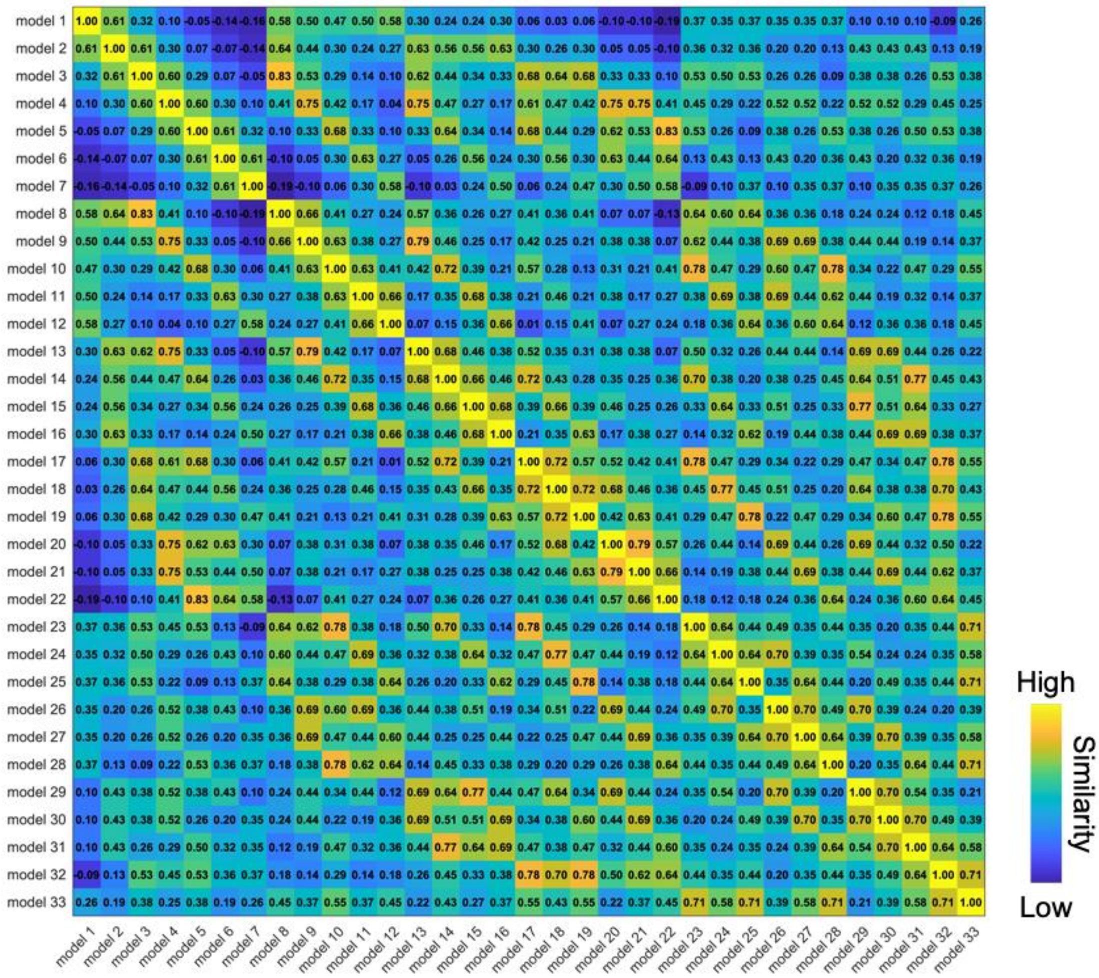

**Figure S 3. Shared variance across theoretical categorical models.** Pairwise Spearman's correlation coefficient across theoretical models of rhythm categorisation. The shared variance reflects the design of the stimuli, which consist in a linear morph between two ratios (here, 1:1 in condition 1 and 2:1 in condition 10).

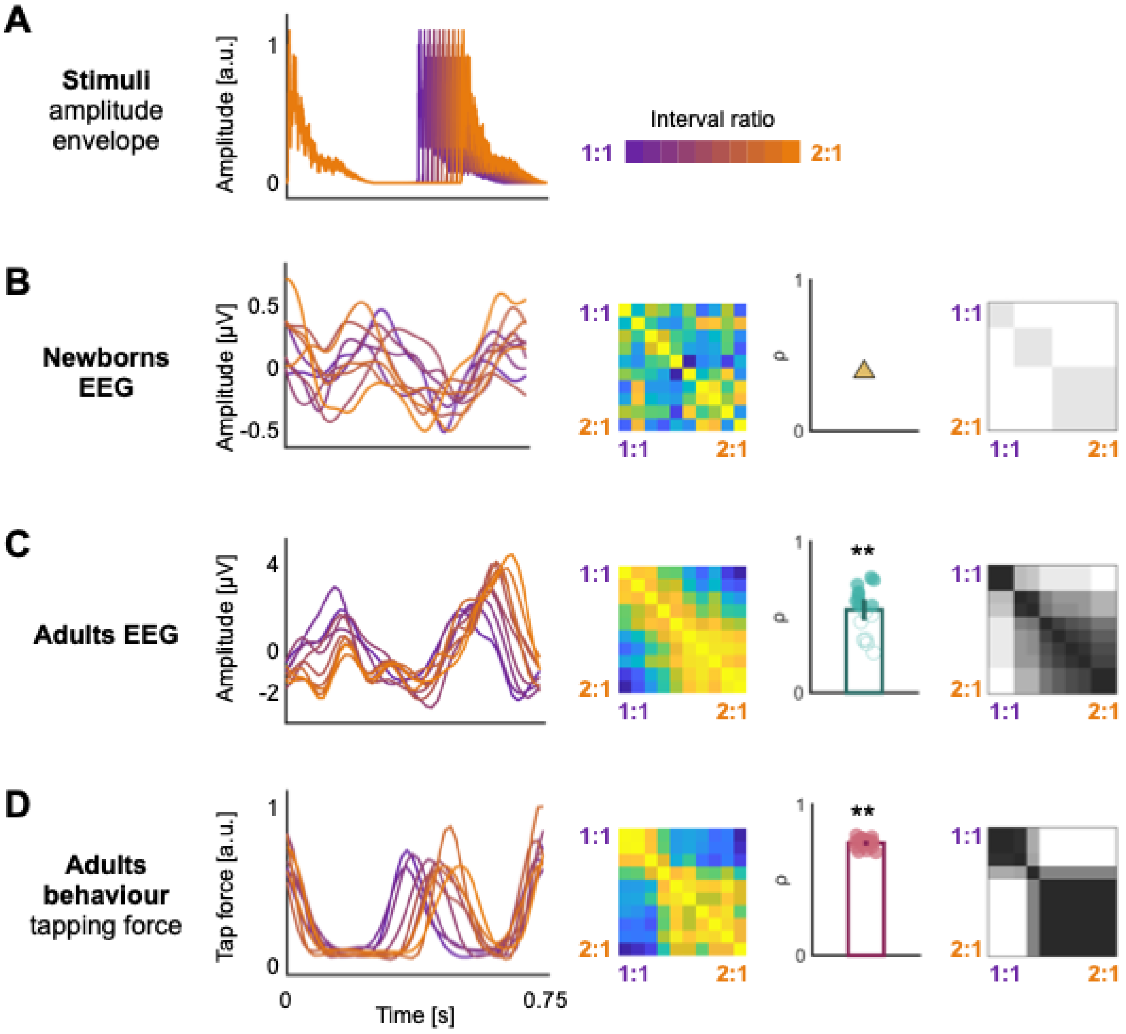


**Figure S 4. Evidence of rhythm categorisation from time domain analyses.** (A) Overlay of one repetition of the rhythmic patterns' envelope across conditions, as indicated by the colour gradient. (B) Time domain results of newborns' neural responses. From left to right: responses over time averaged across participants (fronto-central channels, and pattern repetitions, separately across conditions); RSM built from time domain responses; correlation with the best-fitting theoretical model; best-fitting theoretical model. (C) Time domain results of adults' neural responses. From left to right: responses over time (averaged across fronto-central channels and pattern repetitions) for an example participant; RSM built from time domain responses from an example participant; correlation coefficients with the best-fitting theoretical model (group-level); overlay of best-fitting theoretical models across participants. (D) Same as (C) for tapping responses (continuous tapping force). In all bar plots: the error bars represent the 95% CI, asterisks indicate significant results at the group level (** $p_{Bonf} < 0.01$). In C and D, filled circles indicate significant results at the individual level ($ps_{Bonf} < 0.05$).

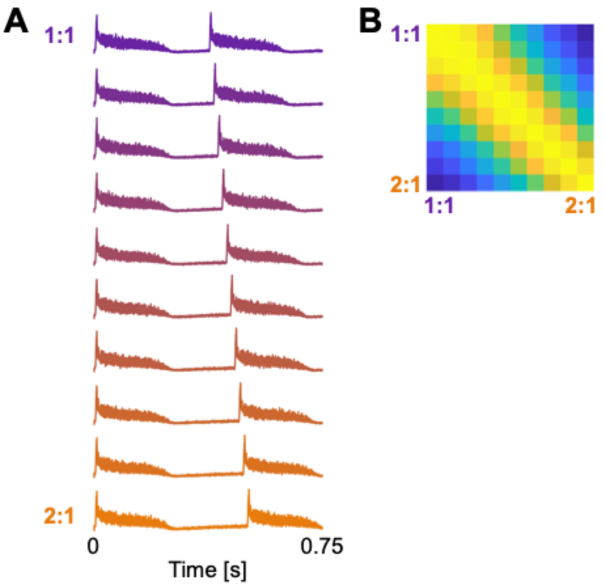


**Figure S 5. Auditory nerve modelled responses and frequency-domain representational geometry.** (A) Modelled auditory nerve responses over a repetition of the rhythmic pattern across conditions (mean spiking rate). (B) Auditory nerve representational similarity matrix (RSM) built from responses in the frequency domain (magnitude spectra) considering frequencies of interest up to 8 Hz (as for the EEG analysis).

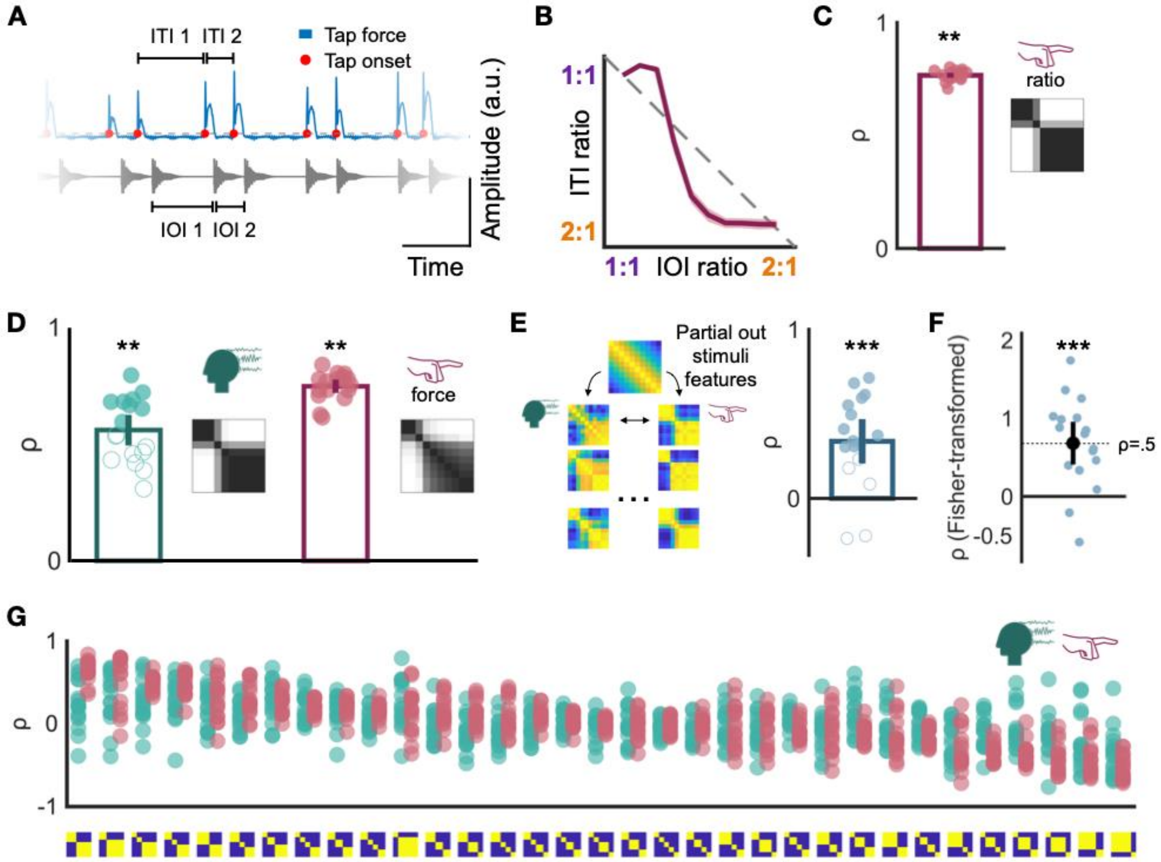


**Figure S 6. Convergence across behavioural and neural categorisation of rhythm in adults.** (A) Excerpt of continuous tapping force with extracted tap onsets (top) to the corresponding rhythmic sequence (bottom). (B) Average ITI ratios as a function of IOI ratios; shaded area corresponds to the standard error of the mean, the diagonal line corresponds to the identity line. (C) Left: correlation of the ITI-ratio RSMs with the best-fitting theoretical model. Right: overlay of best-fitting theoretical model RSMs across participants. (D) Neural results, left panel: correlation of the neural RSMs with the best-fitting theoretical model and overlay of the best-fitting theoretical model RSMs across participants. Behavioural results, right panel: same as for the neural results for tapping force data. (E) Correlation of neural and behavioural RSMs at the individual level after partialling out stimulus RSM. (F) Participant-wise Fisher-transformed correlation between the distribution of correlation coefficients across theoretical models between neural and behavioural responses. (G) Distribution of correlation coefficients across all theoretical models for neural (green) and behavioural (red) responses, ordered by decreasing correlation coefficient of behavioural responses (group average). In all bar plots: the error bar represents the 95% CI, asterisks indicate significant results at the group level (* $p < 0.05$, ** $p < 0.01$, *** $p < 0.001$). In C, D, and E, filled circles indicate significant results at the individual level.

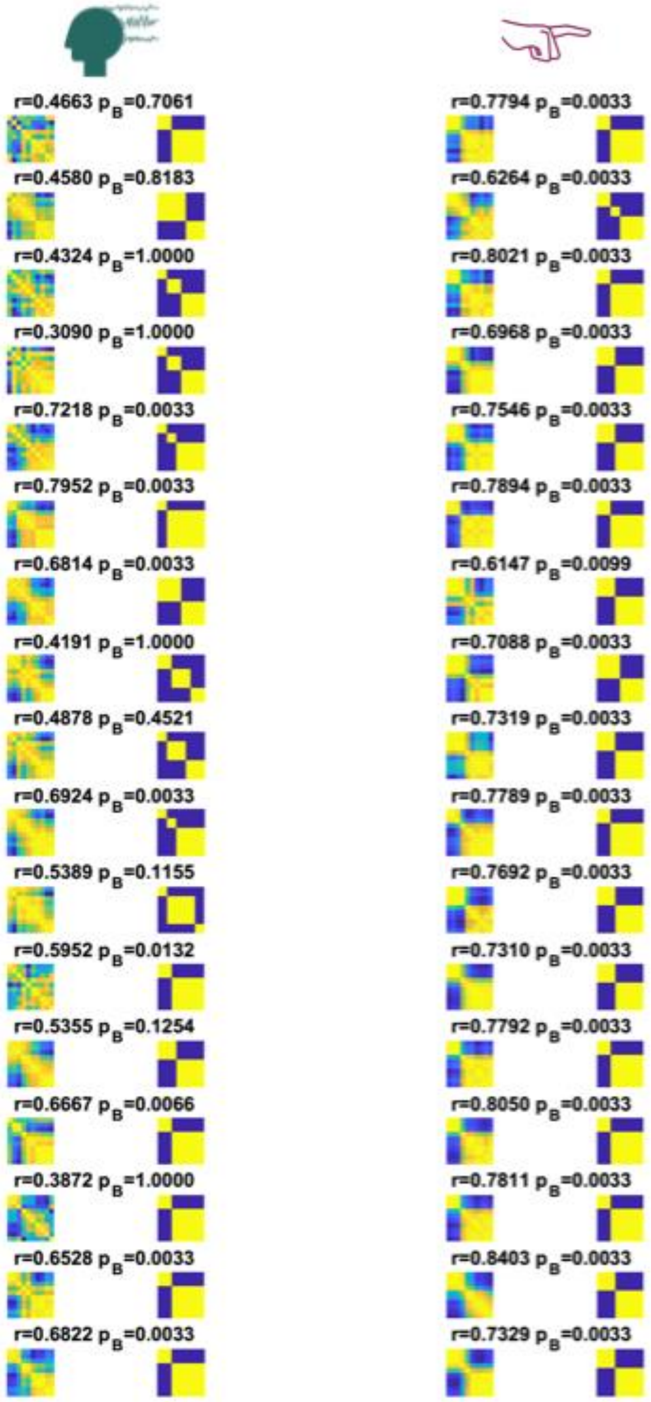


**Figure S 7. Adult individual frequency-RSA results.** Individual representational similarity matrices (RSMs) for neural (EEG, left) and behavioural (tapping force, right) results, together with the corresponding winning theoretical model. Each row corresponds to an individual participant; individual correlation coefficients and p-values (Bonferroni corrected) are reported. For both the continuous tapping force signals and neural signals (corresponding to a pool of 9 fronto-central channels), RSMs were obtained by considering the magnitude spectra up to 8 Hz (first 6 harmonics of the pattern repetition rate).

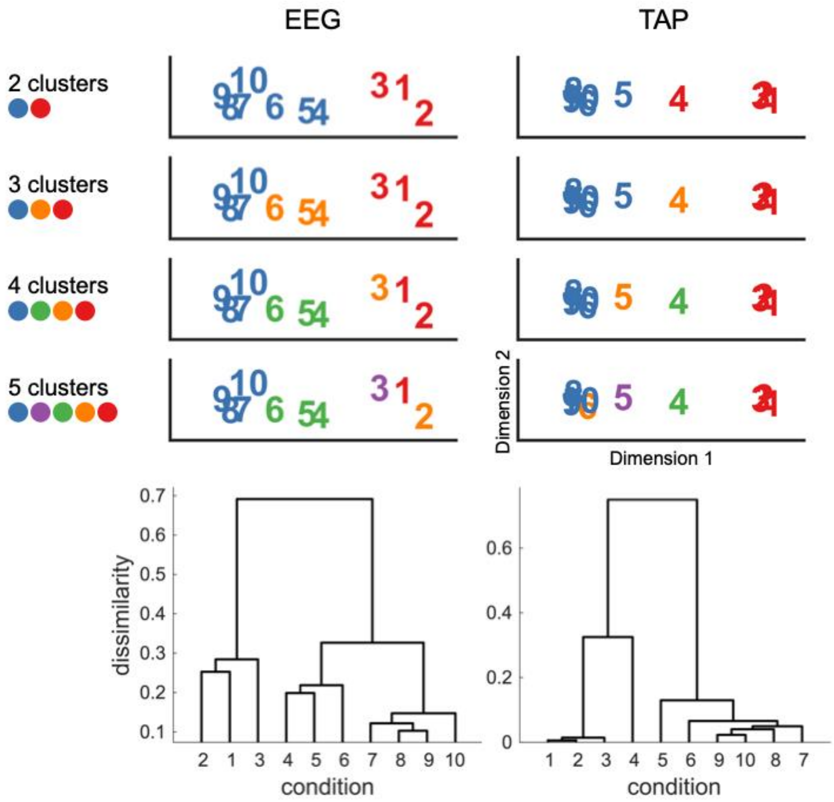


**Figure S 8. Multidimensional scaling and hierarchical clustering of adults' neural and behavioural responses.** Top: hierarchical clustering was implemented on adults' neural (left) and behavioural (right) group-averaged RSMs considering either 2, 3, 4, or 5 clusters (separate rows). In each panel, conditions belonging to the same clusters are visualised with the same colour. Bottom: dendrograms of adults' neural (left) and behavioural (right) responses; the vertical axes represent the dissimilarity distances across conditions. Hierarchical clustering was performed on dissimilarity matrices (1-RSM). In all plots, conditions' numbers from 1 to 10 corresponds to interval ratios from the morphed temporal continuum between 1:1 (condition 1) and 2:1 (condition 10).

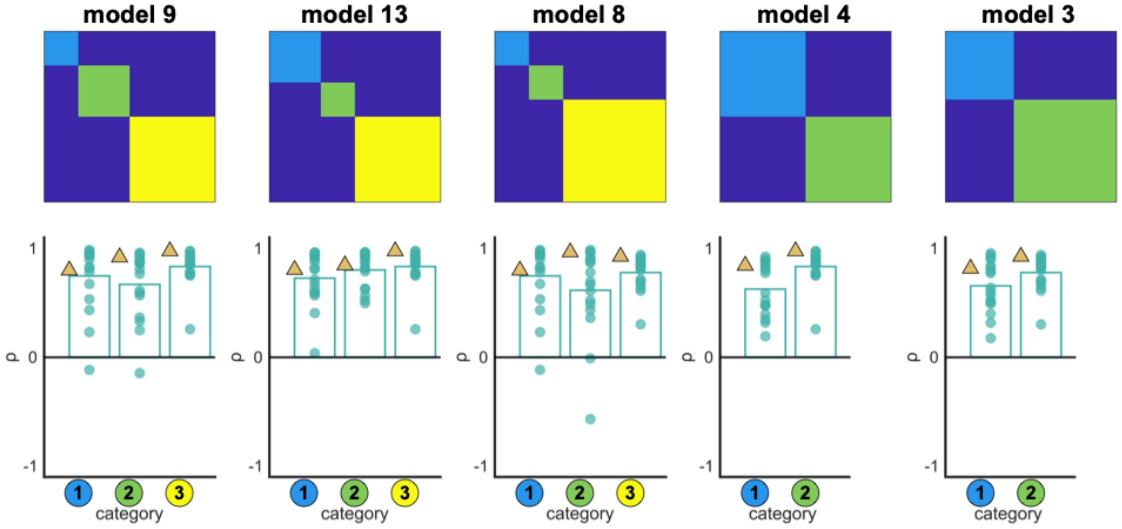


**Figure S 9. Within-category similarity for the theoretical models best explaining newborns' neural data.** For each of the five best-ranked theoretical models describing newborns' neural data (top, ordered from left to right by descending correlation coefficient), we computed the average within-category similarity for newborns' (gold triangle) and individual adult participants' (green circles, bar plot represents group average) neural RSMs. Within-category similarity was computed by averaging the correlation values for each category considering the pixels in the lower triangular part of the RSMs and excluding the diagonal. Qualitatively, these results suggest a higher within-category similarity in the rhythm category on the side of the 2:1 ratio. This is in line with the multidimensional scaling results in newborns, suggesting that conditions closer to 2:1 were more similarly represented than conditions closer to the 1:1 ratio.
